# Susceptibility to inflammatory bowel diseases promotes invasive carcinomas in a murine model of ATF6-driven colon cancer

**DOI:** 10.1101/2024.11.25.624835

**Authors:** Janine Kövilein, Adam Sorbie, Sevana Khaloian, Vanessa Küntzel, Mohamed Ahmed, Sebastian Jarosch, Marianne Remke, Amira Metwaly, Elena M. Reuss, Dirk Busch, Matthieu Allez, Katja Steiger, Barbara Schraml, Olivia I. Coleman, Dirk Haller

**Affiliations:** Technical University of Munich, Chair of Nutrition and Immunology, School of Life Sciences, Freising-Weihenstephan, Germany; Biomedical Center, Ludwig Maximilian University of Munich, Institute for Cardiovascular Physiology and Pathophysiology, Faculty of Medicine, Planegg-Martinsried, Germany; Biomedical Center, Ludwig Maximilian University of Munich, Institute for Immunology, Faculty of Medicine, Planegg-Martinsried, Germany; Institute for Medical Microbiology, Immunology and Hygiene, Technical University of Munich, Munich, Germany; Institute of Pathology, Technical University Munich, Munich, Germany; APHP, Hôpital Saint Louis, Department of Gastroenterology, INSERM UMRS 1160, Paris Diderot, Sorbonne Paris-Cité University, Paris, France; Technical University of Munich, ZIEL – Institute for Food & Health, Freising-Weihenstephan, Germany

**Keywords:** Activating transcription factor 6 (ATF6), ER stress, colitis-associated cancer (CAC), human microbiota associations, IBD-relevant minimal consortium

## Abstract

Chronic inflammation in inflammatory bowel disease (IBD) patients represents a risk factor for developing colitis-associated cancer (CAC). We previously linked the endoplasmic reticulum unfolded protein response (UPR^ER^) signal transducer activating transcription factor 6 (ATF6) with spontaneous microbiota-dependent colonic adenoma development in mice expressing epithelial-specific activated ATF6 (nATF6^IEC^). To investigate IBD-related risk factors in ATF6-mediated tumorigenesis, we crossed tumor-free monoallelic (tg/wt) nATF6^IEC^ mice with Interleukin-10 deficient mice (*Il10*^-/-^). IL10 deficiency initiated tumor susceptibility, with 77% of 12-week tg/wt;*Il10*^-/-^ mice developing colonic adenomas and invasive carcinomas in this novel CAC mouse model. Tumor formation correlated with mucosal immune cell infiltration, characterized by CD11b^+^ granulocytes and monocytes, and mucosa-associated dysbiosis. Colonization of germ-free nATF6^IEC^;*Il10*^-/-^ mice with minimal biosynthetic consortia and IBD stool re-established CAC, confirming microbiota-dependent ATF6-driven tumorigenesis. Increased ATF6 expression in IBD patients during active disease highlights its human relevance. Our findings show that IBD susceptibility heightens the risk for ATF6-driven tumorigenesis.

**Graphical abstract:** 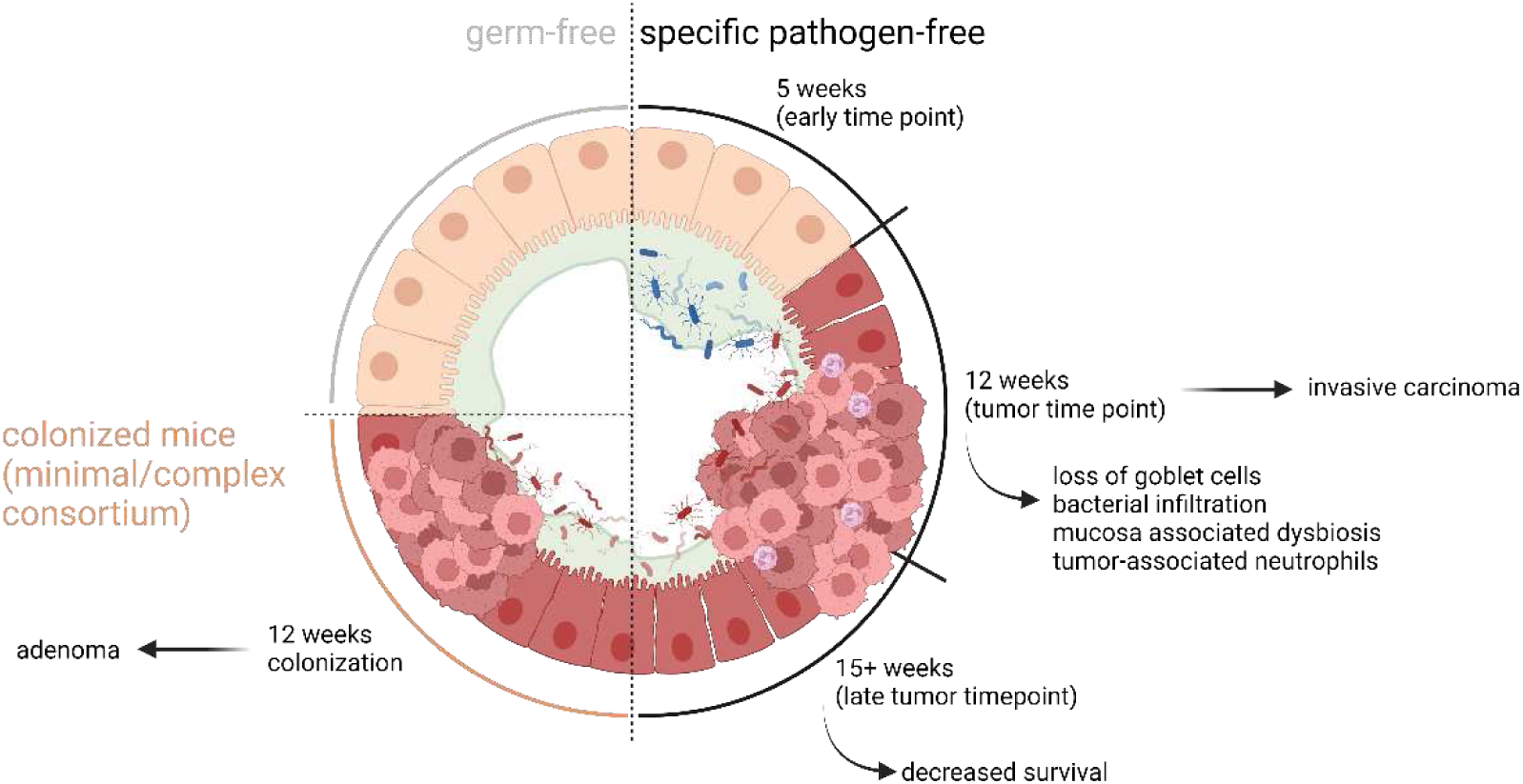

## Introduction

Inflammatory bowel diseases (IBD), including Crohn’s disease (CD) and ulcerative colitis (UC), are highly prevalent diseases with approximately 4.9 million cases worldwide in 2019.^1^ Chronic intestinal inflammation in IBD represents a risk factor for developing colitis-associated cancer (CAC), and induces changes in intestinal microbiota composition. In particular, a decreased abundance of bacteria with putatively beneficial functions, such as *Faecalibacterium prausnitzii*, is observed in IBD and other gastrointestinal (GI) diseases. Consequently, a bloom in pathobionts, such as certain members of Enterobacteriaceae, may occur.^2^ *Escherichia coli* strains encoding the polyketide synthase (*pks*) genomic island, which produce the genotoxin colibactin to cause DNA double-strand breaks,^3^ are commonly found in IBD and colorectal cancer (CRC) patients.^4^ Although a multitude of research has focused on the intestinal microbiome and provides important insights, microbial risk signatures associated with GI diseases remain elusive.^5^ Moreover, most disease-associated taxa have been identified from stool samples rather than biopsies at or near tumor sites, suggesting that such associations likely do not accurately reflect tumor-relevant changes in microbial communities. Indeed, tumor biopsies and corresponding stool samples differ significantly, with fecal microbiota profiles shown to be an unreliable predictor of adenoma formation, suggesting that early alterations in microbiota composition may only be evident at mucosal sites.^6,7^ As such, characterizing mucosal microbial changes occurring before and during tumor onset is important to reliably distinguish between those initiating disease and those promoting progression. To better understand microbe-host interactions in the context of tumorigenesis, we recently generated a transgenic mouse model of intestinal epithelial cell (IEC)-specific expression of the active p50 fragment of the endoplasmic reticulum (ER) transmembrane protein activating transcription factor 6 (nATF6), representing a mouse model for spontaneous CRC.^8^ In eukaryotic cells, the ER is responsible for correct protein folding to maintain intestinal tissue homeostasis. Adequate protein folding in the ER is orchestrated by a group of conserved signaling pathways, collectively termed the unfolded protein responses (UPR^ER^), with ATF6 comprising one of those pathways. Under normal conditions, Glucose-regulated protein 78 (Grp78) is bound to ATF6, keeping the UPR^ER^ in an inactive state. The accumulation of un- or misfolded proteins leads to the dissociation of Grp78 from the UPR^ER^ transmembrane proteins, starting signaling cascades aiming at restoring proteostasis or promoting apoptosis.^9^ However, chronic UPR^ER^ activation or ER-stress can lead to inflammation and cancer.^9,10^ Recently, we identified a strong link for a functionally and genetically impaired interaction between the UPR and autophagy in IBD, with ATF6 increasing pro-inflammatory cytokine expression in response to ER stress *in vitro*.^11^

We previously showed that ATF6 signaling provides a novel link between the UPR^ER^ and spontaneous CRC in mice.^8^ Biallelic activation of nATF6 in murine IECs resulted in spontaneous intestinal tumorigenesis, with a loss of mucin-filled goblet cells and increased microbial penetration to the epithelium preceding tumor onset. While tumorigenesis was microbiota-dependent, the development of inflammation was the consequence rather than the cause of tumor formation. We showed that a subset of patients of The Cancer Genome Atlas (TCGA) dataset show genomic and transcriptomic aberrations in ATF6, underlining the human relevance of our model.^8^ Moreover, ATF6 was proposed as a marker for early dysplastic changes in CAC and CRC, with high ATF6 expression predicting neoplastic transformation with high accuracy.^12^ In this study, we characterize the impact of IBD-related risk factors on ATF6-mediated tumorigenesis in our newly generated mouse model. By crossing nATF6^IEC^ mice with Interleukin-10 deficient mice (*Il*10^-/-^) and the transfer of IBD-relevant microbiota, we created an inflammatory background mimicking CAC. Interestingly, IL10 deficiency increased ATF6-dependent tumor susceptibility and caused the formation of invasive carcinomas in a microbiota-dependent manner.

## Results

### Interleukin-10 deficiency increases tumor susceptibility in nATF6^IEC^ mice and drives formation of colonic invasive carcinomas

We previously showed that biallelic (tg/tg), but not monoallelic (tg/wt) nATF6^IEC^ mice develop early-onset colonic adenomas in a microbiota-dependent manner by the age of 12 weeks.^8^ Inflammation was shown to be a consequence rather than the cause of ATF6-mediated tumorigenesis.^8^ To test the hypothesis of whether IBD susceptibility affects tumor onset and phenotype, we crossed biallelic and monoallelic nATF6^IEC^ mice with *Il10^-/-^* mice. IL10 deficiency in biallelic nATF6^IEC^ mice (tg/tg;*Il10^-/-^*) substantially accelerated disease onset, and mice did not exceed the age of 10 weeks (Fig. 1A). Interestingly, the combination of the monoallelic nATF6^IEC^ model with IL10 deficient mice (tg/wt;*Il10^-/-^*) showed markedly decreased survival, reaching similar mortality levels when compared to the biallelic nATF6^IEC^ model (Fig. 1A). Quantitative real-time PCR (qRT-PCR) measurements and immunohistochemical (IHC) staining of ATF6 expression and its downstream target *Grp78* confirmed activation and downstream signaling of ATF6 in tg/wt;*Il10^-/-^* mice across all time points (5 weeks = pre-tumor, 12 weeks = tumor and 15+ weeks = late tumor time point) in contrast to fl/fl;*Il10^-/-^* controls (supplemental Fig. 1A and 1B). Importantly, IL10 deficiency clearly increased tumor susceptibility, with 77% and 61% of tg/wt;*Il10^-/-^* mice displaying tumors at 12 weeks and 15+ weeks, respectively, while fl/fl;*Il10^-/-^* controls stayed tumor-free (NT; non-tumor) (Fig. 1B). Tumor number and volume in tg/wt;*Il10^-/-^* mice were similar at both tumor time points (Fig. 1C and supplemental Fig. 1C). In line with our previous findings,^8^ tumor formation was restricted to the proximal to mid part of the colon, revealing that inflammation does not alter the spatial localization of tumorigenesis in our murine model. Since not all tg/wt;*Il10^-/-^* mice developed tumors, mice were categorized into responders (R; mice with tumors) and non-responders (NR; mice without tumors, Fig. 1B). Histological examination and tumor scoring of the tumor susceptible (proximal) and the non-susceptible (distal) regions of colon Swiss rolls revealed substantial variation of tumor phenotypes in tg/wt;*Il10^-/-^* mice. Strikingly, while dysplasia scores were similar at both tumor time points in the susceptible and non-susceptible colon regions (Fig. 1D and supplemental Fig. 1D), we observed the development of invasive carcinomas in a subset of R tg/wt;*Il10^-/-^* mice (dysplasia score 4, Fig. 1D and 1E). Through a spatially resolved analysis, in which the colon was separated into 0.5 cm segments, we could classify the colon into a tumor susceptible region (colon tissue site 1 to 9) and a non-susceptible region (colon tissue site 10 onwards, Fig. 1F). Tumor incidences were highest in the first two tissue sites and decreased towards the mid colon (Fig. 1F). To better characterize the tumorigenic phenotype, we histologically quantified proliferating cells (anti-Ki67) and mucin-filled goblet cells (GCs, Periodic Acid-Schiff Alcian Blue (PAS-AB)) in the tumor susceptible colon region of 15+ week old mice. Proliferation was significantly higher in tg/wt;*Il10^-/-^* mice compared to fl/fl;*Il10^-/-^* with highest expression in tumor regions of R tg/wt;*Il10^-/-^* mice (Fig. 1G). We next analyzed GCs, as GCs are secretory cells in the intestinal tract that require a homeostatic environment in the ER to function correctly and loss of mucin-filled GCs was a key characteristic of our nATF6^IEC^ mouse model preceding tumor development.^8^ Here, tg/wt;*Il10^-/-^* mice showed a decrease in mucin-filled GCs compared to fl/fl;*Il10^-/-^* mice, with selected crypts in tumor regions showing a complete loss of mucin-filled GCs (Fig. 1G).

**Fig. 1.**
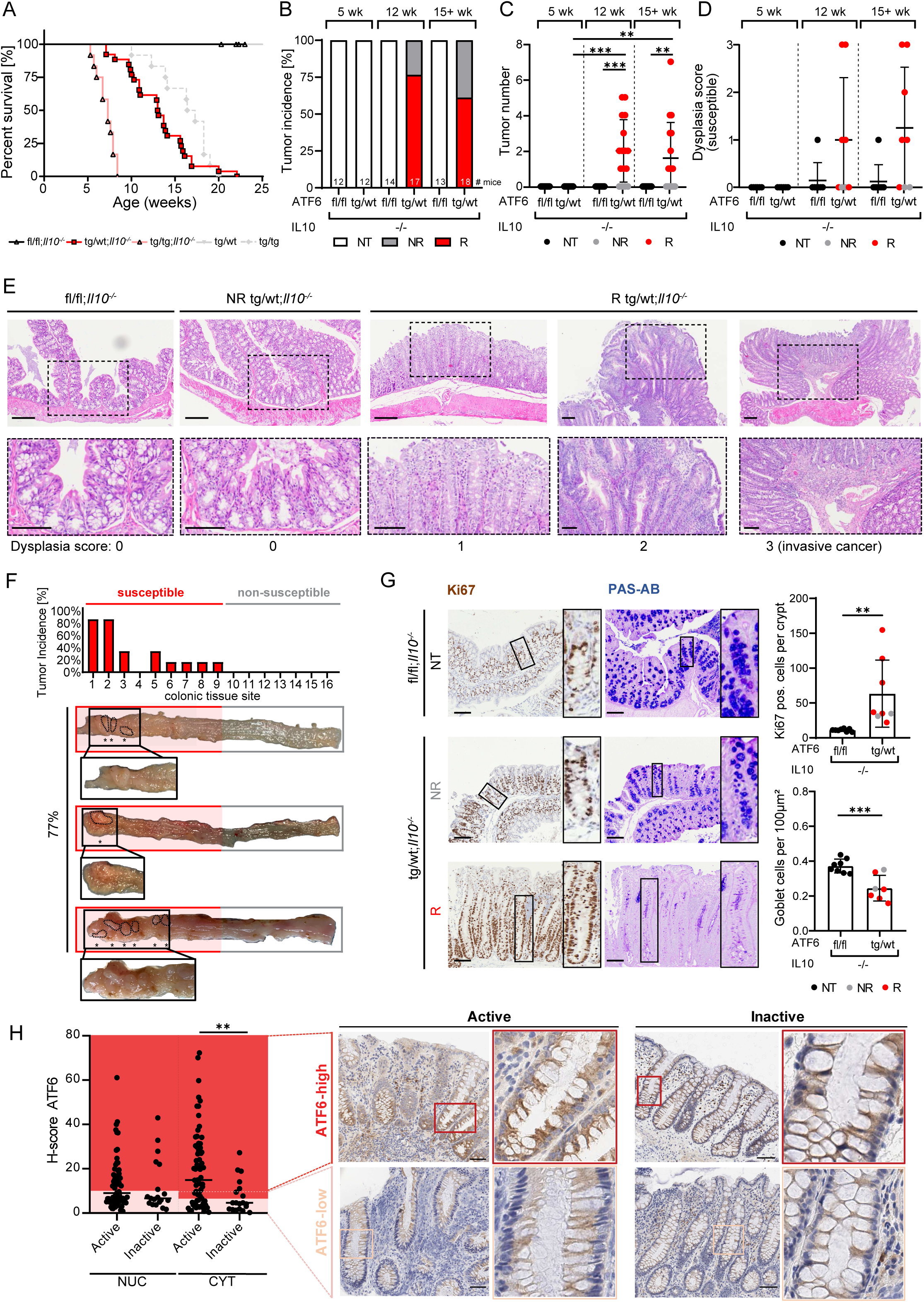
Interleukin-10 deficiency increases tumor susceptibility in nATF6^IEC^ mice and drives formation of colonic invasive carcinomas. **A)** Survival curves of nATF6^IEC^;*Il10^-/-^* mice housed under specific pathogen-free (SPF) conditions. **B)** Colonic tumor incidence of tg/wt;*Il10^-/-^* mice categorized into responder (R = mice that developed tumors) and non-responder (NR = mice that did not show any tumors) at week 5, 12 and 15+. **C)** Colonic tumor numbers of nATF6^IEC^;*Il10^-/-^* mice at week 5, 12, and 15+. **D)** Dysplasia score resulting from histologically scoring of susceptible (proximal) colon tissue of nATF6^IEC^;*Il10^-/-^* mice for mucosal architecture and atypia (0-3) at week 5, 12, and 15+. **E)** Representative images of H&E staining of susceptible colon tissue at week 12 (scale bars: 200 µm) and corresponding higher magnifications (*rectangles*; scale bars: 100 µm). **F)** Tumor incidence of 0.5 x 0.5 cm colon tissue sites identifying a tumor susceptible (colon tissue site 1 to 9) and non-susceptible (colon tissue site from 10 onwards) region with representative macroscopic images of tg/wt;*Il10^-/-^* mice; asterisks indicate tumors. **G)** Ki67 and PAS-AB staining of proximal or susceptible colon tissue of fl/fl;*Il10^-/-^* and NR and R tg/wt;*Il10^-/-^* mice (scale bars: 100 µm) and corresponding higher magnifications (*rectangles*; scale bars: 100 µm). Respective quantifications of the staining were analyzed in tumor susceptible colon tissue. **H)** H-score of nuclear (NUC) and cytoplasmic (CYT) ATF6 expression in colon biopsies of IBD patients stained immunohistochemically. Representative images of ATF6 expression in active and inactive disease as well as ATF6-high and ATF6-low expression in colon tissue (scale bars: 100 µm) and corresponding higher magnifications. wk = week; NT = non-tumor; NR = non-responder; R = responder; NUC = nuclear; CYT = cytoplasmic.

To investigate the relevance of ATF6 in IBD, we next analyzed an IBD cohort (CD n=36, UC n=32; supplemental Table 1) in which patients were followed for one year after initiation of therapy, to assess the dynamics of stool and mucosa-associated gut microbiota composition (three sampling time points). To validate our findings in a human setting, we quantified ATF6 protein expression using IHC staining in a subset of the afore-mentioned cohort (n=64). Nuclear (NUC) as well as cytoplasmic (CYT) ATF6 was quantified using QuPath, revealing a significantly increased CYT ATF6 expression during active disease compared to inactive disease (Fig. 1H).

Taken together, these data clearly suggest that inflammation accelerates ATF6-driven colonic tumorigenesis with otherwise tumor-free tg/wt mice developing invasive carcinomas when crossed with IL10 deficient mice. Furthermore, we demonstrate that IBD patients with active disease show increased ATF6 expression, confirming clinical relevance.

### IL-10 deficiency triggers innate immune cell infiltration into the mucosa of monoallelic nATF6^IEC^ mice

To understand the contribution of immune cells in the different colon regions, we performed immune characterization of tumor susceptible and non-susceptible colon tissue of nATF6^IEC^;*Il10^-/-^* mice. Inflammation scores were similar in both the tumor susceptible and non-susceptible colon region in tg/wt;*Il10^-/-^* mice compared to fl/fl;*Il10^-/-^* controls at all three time points (Fig. 2A). Lipocalin-2 (LCN2) was measured in fecal samples, and showed higher levels in tg/wt;*Il10^-/-^* mice (both R and NR) compared to fl/fl;*Il10^-/-^* control mice (Fig. 2B), confirming an increased inflammatory response in tg/wt;*Il10^-/-^* mice independent of tumor formation. To identify the different immune cell populations in tumor susceptible and non-susceptible colon regions of nATF6^IEC^;*Il10^-/-^* mice at the onset of tumor formation using R tg/wt;*Il10^-/-^* mice, immune cells were isolated from the lamina propria and characterized by spectral flow cytometry. Different Uniform Manifold Approximation and Projection (UMAP) were generated either on CD3^+^CD90^+^ lymphocytes or on remaining CD3^-^CD90.2^-^ cells. Nine and eight clusters were identified within the T cells and ILCS (CD3^+^CD90.2^+^) and the CD3^-^CD90^-^ cells, respectively (Fig. 2C). R tg/wt;*Il10^-/-^* mice showed an increase in overall leukocyte cell count per square (0.5 cm^2^ colon tissue) compared to fl/fl;*Il10^-/-^* control mice. Interestingly, the number of leukocytes was highest in the non-susceptible tumor-free colon tissue of R tg/wt;*Il10^-/-^* mice (Fig. 2D and E). Notably, the frequency and number of neutrophils (CD11b^+^Ly6G^+^cells) and monocytes (CD11b^+^Ly6C^+^ cells) per square were increased in R tg/wt;*Il10^-/-^* mice compared to fl/fl;*Il10^-/-^* controls, especially in non-susceptible tumor-free regions (Fig. 2E). CD4^+^ T cells were twice as frequent as neutrophils and monocytes, and showed similar percentages in both genotypes. The increased frequency of CD4^+^ T cells in tumor susceptible colon regions can be attributed to higher percentages of gut-specific RORγt^+^Tregs (CD4^+^Foxp3^+^RORγt^+^) in the same tumor susceptible region (Fig. 2E and supplemental Fig. 2A). A similar trend was observed for classical T_regs_ (CD4^+^Foxp3^+^; supplemental Fig. 2A). As these differences were found in both genotypes, they are likely inflammation rather than tumor-driven.

**Fig. 2.**
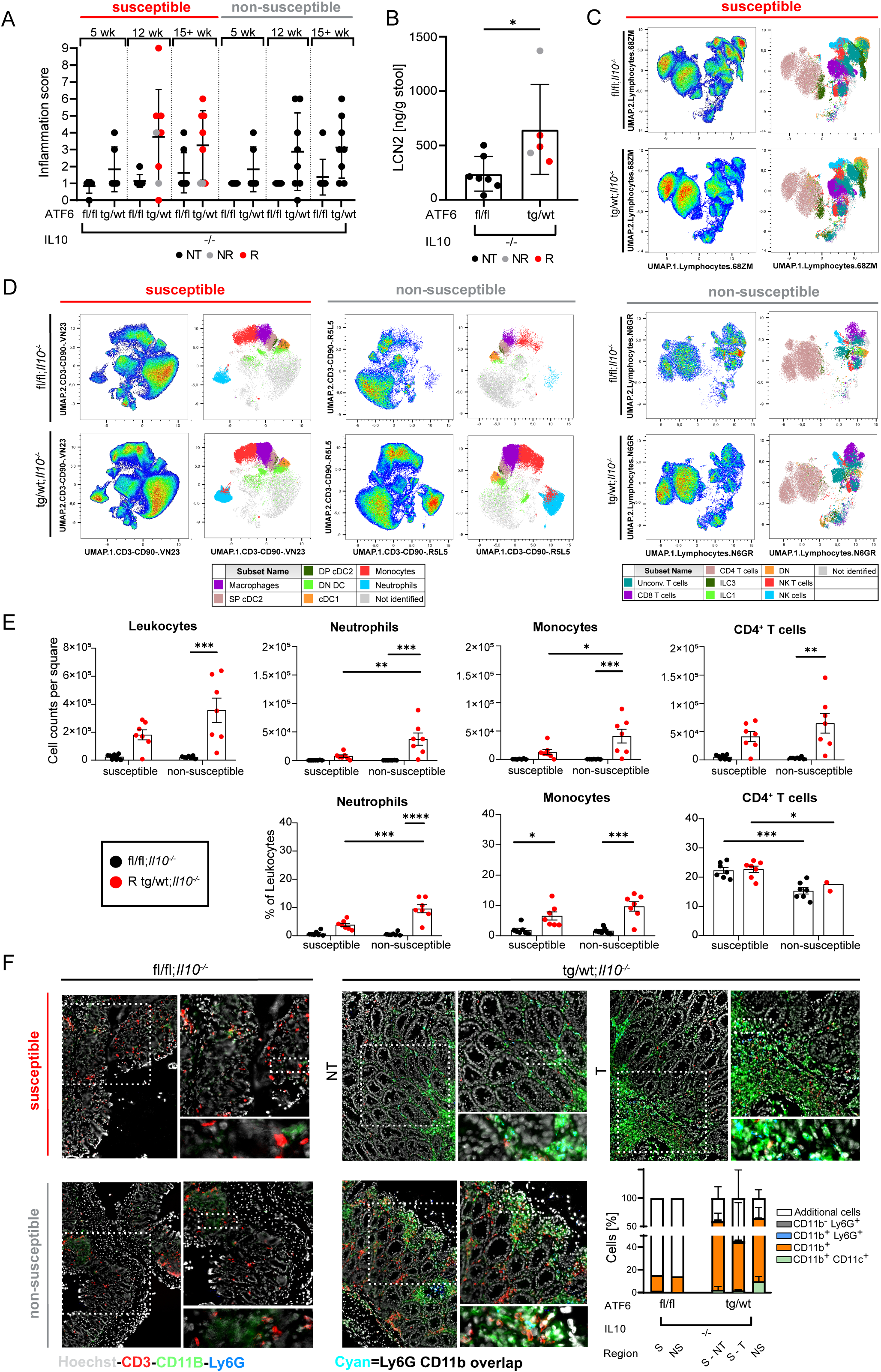
nATF6^IEC^;*Il10^-/-^* mice show changes in innate immunity. **A**) Inflammation score for susceptible (proximal) and non-susceptible (distal) colon tissue. nATF6^IEC^;*Il10^-/-^* mice were histologically scored for immune cell infiltration and epithelial damage resulting in an inflammation score (0-9). **B)** Lipocalin-2 (LCN2) levels in fecal samples of nATF6^IEC^;*Il10^-/-^* mice. **C)** UMAPs depicting cell abundance and distribution of CD3^+^CD90^+^ lymphocytes in colon tissue of nATF6^IEC^;*Il10^-/-^* mice. Colon tissue was divided according to tumor susceptible region (proximal-mid = 2/3 colon length) and non-susceptible region (distal = 1/3 colon length). **D)** UMAPs depicting cell abundance and distribution of remaining CD3^-^CD90.2^-^ cells of susceptible and non-susceptible colon tissue of nATF6^IEC^;*Il10^-/-^* mice. **E)** Quantification of total leukocytes, neutrophils, monocytes and CD4^+^ T cells given as cell counts per square (0.5 x 0.5 cm colon tissue) and % of leukocytes. **F)** Quantification of immune cell populations in the colonic lamina propria of R nATF6^IEC^;*Il10^-/-^* mice at 15+ weeks of age by ChipCytometry staining. Immune cells are stained as follows: CD3 (red), CD11b (green), Ly6G (blue). Nuclei are stained with Hoechst (grey). R = responder; NT = non-tumor; T = tumor.

The frequency of ILC3s and NK cells decreased in the susceptible region of R tg/wt;*Il10*^-/-^ mice compared to fl/fl;*Il10^-/-^* controls (supplemental Fig. 2A). A similar drop in frequencies in the non-susceptible region suggests that observed alterations may contribute to but are not solely driving local tumor formation. CD8^+^ cells (supplemental Fig. 2A) and CD8^+^ memory and effector subsets did not show any tumor-relevant alterations (data not shown). Unconventional T cells (CD4^-^CD8^-^) showed a reduction in the frequency in R tg/wt;*Il10*^-/-^ mice compared to fl/fl;*Il10^-/-^* controls (supplemental Fig. 2A), in both the susceptible as well as non-susceptible colon regions, pointing towards a global genotype rather than tumor-phenotype effect. Similarly, alterations in NK T cell frequency (CD3^+^NK1.1^+^) suggest colonic region-specific alterations independent of ATF6-driven tumor relevance (supplemental Fig. 2A). Overall, analyzed immune cell frequencies shift towards a more myeloid-based immune composition.

We next used ChipCytometry to specifically dissect immune infiltration in tumor and non-tumor tissue within the tumor susceptible colon region. In line with our spectral flow cytometry findings, an overall increase in CD11b^+^ cells was observed in tg/wt;*Il10^-/-^* mice compared to fl/fl;*Il10^-/-^* mice (Fig. 2F). Furthermore, more Ly6G^+^ cells were detected in tg/wt;*Il10^-/-^* mice, especially in the tumor compartment, compared to fl/fl;*Il10^-/-^* controls (Fig. 2F and supplemental Fig. 2A). An overlapping signal of Ly6G and CD11b (depicted in cyan) was present mainly in tumor regions of tg/wt;*Il10^-/-^* mice, validating the above finding in R tg/wt;*Il10^-/-^* mice and confirming that neutrophils specifically infiltrate the tumor tissue.

Together, these data suggest an overall increased inflammatory response in tg/wt;*Il10^-/-^* mice, showing that abnormalities in innate immune cells, specifically the infiltration of tumor-associated neutrophils, constitute a key characteristic of the tumor susceptible region.

### The mucosa-associated microbiota of responder mice is enriched in CAC-relevant pathobionts

We previously demonstrated that tumor development in nATF6^IEC^ mice is microbiota dependent.^8^ We therefore sought to investigate microbial changes occurring at the mucosal and luminal (cecal) sites in nATF6^IEC^;*Il10^-/-^* mice. Profiling of mucosa-associated microbiota at tumor onset revealed that various measures of α-diversity, including richness (number of species), Shannon effective (evenness) as well as Faith’s PD (phylogenetic diversity) were decreased in R tg/wt;*Il10^-/-^* mice compared to both NR tg/wt;*Il10^-/-^* mice and fl/fl;*Il10^-/-^* controls (Fig. 3A). In line with these findings, mucosal β-diversity, based on generalized UniFrac distance^13^, showed microbial profiles of R tg/wt;*Il10^-/-^* mice to be distinct from NR tg/wt;*Il10^-/-^* mice (Fig. 3B). Interestingly, analysis of the same α- and β-diversity metrics in luminal microbiota profiles did not reveal any significant differences (Fig. 3A and 3B). To substantiate these observations, we trained multiple machine learning models on both mucosal and luminal data and examined their ability to discriminate between the T and NT phenotypes. Across all models tested, mucosal data outperformed luminal data, indicating an inherent tumor-associated signature present in mucosal data, which is absent in luminal samples (Fig. 3C).

**Fig. 3.**
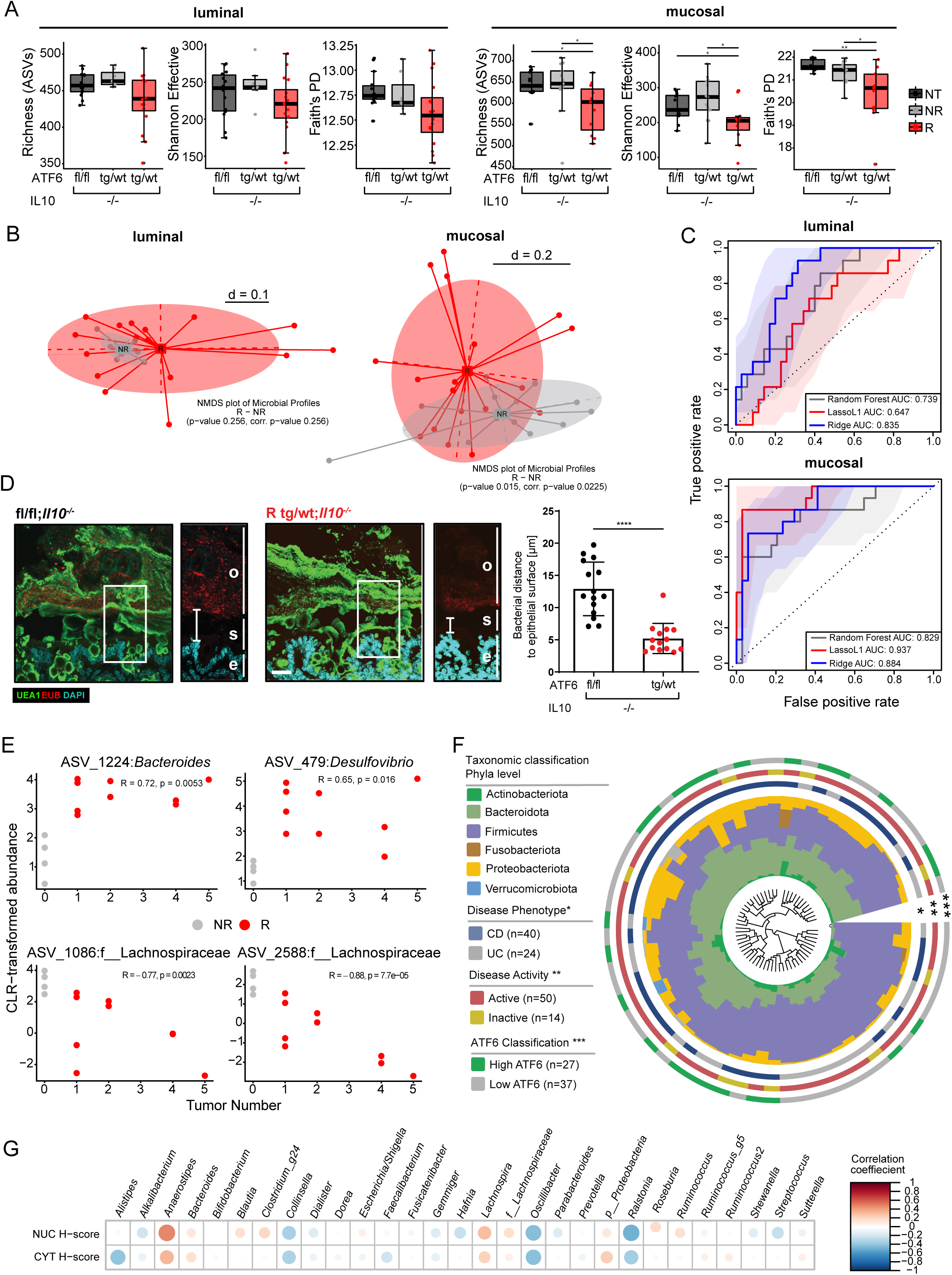
Mucosa-associated microbiota of responder mice is enriched in CAC-relevant pathobionts. **A)** Luminal (cecal) and mucosal (colonic) α-diversity (richness, Shannon Effective and Faith’s phylogenetic diversity) in fl/fl;*Il10^-/-^* control mice and NR and R tg/wt*;Il10^-/-^* mice. **B)** Luminal and mucosal β-diversity between NR and R tg/wt;*Il10^-/-^* mice based on generalized UniFrac distance. Differences between groups were tested using PERMANOVA. **C)** ROC curves comparing true vs false positive rates of Random Forest, L1-penalised Lasso and Ridge regression models trained on luminal and mucosal data. Mucosal data was randomly subsampled to match luminal sample size (original n=57, subsampled=49). All models were trained using repeated 5-fold cross-validation with 5 repeats. **D)** Fluorescent *in situ* hybridization (FISH) targeting 16S rRNA (EUB338 probe) in combination with immunostaining of mucus (UEA1) in the proximal colon of fl/fl;*Il10^-/-^* and tg/wt;*Il10^-/-^* mice (scale bars: 100 µm) and corresponding higher magnifications (*rectangles*; scale bars: 100 µm). Nuclei are counterstained with DAPI. Quantification of the bacterial distance to the epithelial surface.o = outer layer, s = stratified layer, e = epithelia. **E)** Spearman-correlation of centered-log-ratio (CLR) transformed abundance of tumor enriched ASVs classified as Bacteroides, Desulfovibrio and Lachnospiraceae and tumor number in 12-week-old tg/wt;*Il10^-/-^* mice. **F)** Phylogenetic tree showing the similarities between microbiota profiles based on generalized UniFrac distances in mucosal biopsies derived from CD patients (n=40) and UC patients (n=24). Individual taxonomic composition at the phylum level is shown as stacked bar plots around the phylogram. Innermost ring shows stratification based on disease phenotype, CD patient samples (blue) and UC patient samples (grey) and denoted by an asterisk (*); the second ring shows stratification based on disease activity, active disease (red), inactive disease (yellow) and denoted by two asterisks (**); the third ring shows stratification based on ATF6 classification, high ATF6 (green), low ATF6 (grey) and denoted by three asterisks (***). **G)** Correlations of H-Scores of nuclear (NUC) and cytoplasmic (CYT) ATF6 expression in colon biopsies of IBD patients with bacterial genera. NT = non-tumor; NR = non-responder; R = responder; AUC = area under the curve; o = outer layer, s = stratified layer, e = epithelia; UEA1 = ulex europaeus agglutinin 1; EUB = Eubacteria FISH probe; DAPI = 4’,6-diamidino-2-phenylindole; ASV = amplicon sequence variant; CD = Crohn’s disease; UC = ulcerative colitis; NUC = nuclear; CYT = cytoplasmic.

Since a dysfunctional host mucus layer often precedes inflammatory and tumorigenic diseases, and can cause changes in the composition of the mucosal microbiota, we next examined the structural integrity of the mucus layer in nATF6^IEC^;*Il10^-/-^* mice. Combined visualization of mucosal-resident microbes using Fluorescence in situ hybridization (FISH), and the mucus layer, via anti-Ulex Europeaus Agglutinin 1 (UEA1) fluorescence lectin staining of Carnoy-fixed colon tissue, revealed a decreased bacterial distance to the epithelial surface in tg/wt;*Il10^-/-^* mice compared to fl/fl;*Il10^-/-^* controls, as well as a structural loss of the stratified inner mucus layer (Fig. 3D). Differential abundance analysis of mucosal data identified several taxa enriched in R mice, mostly comprising ASVs classified as Mucispirillum, Bacteroides and Desulfovibrio, while depleted taxa were mostly represented by Lachnospiraceae (supplemental Fig. 3A and 3B). Spearman-rank correlation of centered log-ratio (CLR) transformed abundance between those ASVs and tumor number in R tg/wt;*Il10^-/-^* mice revealed that Bacteroides and Desulfovibrio correlated positively with tumor numbers, while two Lachnospiraceae ASVs correlated negatively (Fig. 3E). Mucispirillum however, did not correlate with tumor number (supplemental Fig. 3D).

Lastly, we performed 16S rRNA amplicon sequencing of the mucosa-associated microbiota in the above-described IBD cohort, to verify our findings. Stool microbiota data analysis was previously published by Urbauer *et al*. ^14^ Phylogenetic tree based on generalized UniFrac distances in mucosal biopsies showed no separation of the microbiota profiles of CD (n=79) and UC (n=70) mucosal samples (supplement Fig. 3E). Taxonomic classification on phylum level showed prevalence of Bacteroidota, Firmicutes and Proteobacteriota. Most of the biopsies were sampled during active disease (n=119 *vs*. n=30 inactive disease). α-diversity, including richness (number of species) and Shannon effective, were increased in UC patients during both active (A) and inactive (IN) disease (supplemental Fig. 3F). β-diversity showed microbial profiles of CD patients being distinct from UC patients (supplemental Fig. 3G), with ASVs classified as Bacteroides being enriched in CD patients, similar to our murine model (supplemental Fig. 3H). Cohort stratification based on disease phenotype, disease activity and ATF6 classification showed no differences at phylum level in the subset of the cohort (Fig. 3F). Interestingly, however, high ATF6 expression correlated with an increased CLR-transformed abundance of several of the same bacterial genera that were previously found to be increased in our murine model (e.g. Bacteroides and Lachnospira; Fig. 3G). Taken together, mucosa-associated microbiota better predicts the tumorigenic phenotype compared to the luminal microbiota, with alterations in microbial composition restricted to R tg/wt;*Il10^-/-^* mice.

### IBD-relevant bacterial consortia and complex microbiota from IBD patients re-established tumor formation in germ-free nATF6^IEC^;*Il*10^-/-^ mice

To verify whether the absence of bacteria prevents tumor formation in our CAC model in the same way as previously shown in our CRC model,^7^ nATF6^IEC^;*Il10^-/-^* mice were housed under GF conditions. Indeed, we confirmed the absence of tumor formation in GF mice sampled at the age of 25 weeks. Furthermore, GF mice showed no reduced survival, in contrast to SPF mice (Fig. 4A and 1A). Histopathological scoring of GF fl/fl;*Il10^-/-^* and tg/wt;*Il10^-/-^* mice showed no signs of dysplasia and no change in epithelial damage or immune cell infiltration (Fig. 4B-F). To investigate whether an IBD-relevant consortium enhances tumorigenesis, we colonized nATF6^IEC^;*Il10^-/-^* mice with an IBD-relevant minimal biosynthetic consortium (modified version of simplified human microbiota, SIHUMI).^15^ Mice were colonized at the age of four weeks for a period of 12 weeks, after which tumor formation would be expected. The modified SIHUMI used in this study consisted of six bacterial strains (*Enterococcus faecalis*, *Ruminococcus gnavus*, *Phocaeicola vulgatus* (formerly *Bacteroides vulgatus*), *Lactobacillus plantarum*, *Bifidobacterium longum* and the disease relevant *Escherichia coli* strain (murine derived *pks*^+^ NC101)).^4^ Interestingly, colonized tg/wt;*Il10^-/-^* mice already showed reduced survival compared to colonized fl/fl;*Il10^-/-^* and GF mice six weeks after gavage (Fig. 4A). Tumor formation occurred in SIHUMI colonized tg/wt;*Il10^-/-^* mice, with tumor incidence (64.3%; Fig. 4B) and tumor numbers (up to 7 tumors; Fig. 4C) comparable to SPF tg/wt;*Il10^-/-^* mice at 12 weeks of age (Fig. 1B and 1C). Tumor volume, however, was clearly lower in SIHUMI colonized tg/wt;*Il10^-/-^* mice (up to 17 mm^3^) compared to SPF tg/wt;*Il10^-/-^* mice (up to 75 mm^3^; Fig. 4D and supplement Fig. 1C). Although dysplasia scores were increased in SIHUMI colonized tg/wt;*Il10^-/-^* mice in the susceptible and non-susceptible colon regions (Fig. 4E and supplemental Fig. 4A, B), none of the tumors in tg/wt;*Il10^-/-^* mice were classified as invasive carcinomas, in contrast to SPF tg/wt;*Il10^-/-^* mice (Fig. 1D). Tumors in SIHUMI colonized tg/wt;*Il10^-/-^* mice were classified as adenomas (dysplasia score 2), as previously shown in biallelic nATF6^IEC^ mice.^7^ In contrast, inflammation scores of SIHUMI colonized tg/wt;*Il10^-/-^* mice did reach levels similar to those observed in SPF mice, in all colon regions (Fig. 4F and supplement Fig. 4C). Furthermore, LCN2 was enriched in both R and NR tg/wt;*Il10^-/-^* mice compared to fl/fl;*Il10^-/-^* control mice and GF mice, confirming the presence of inflammation (Fig. 4G). Proliferation in SIHUMI colonized mice was enhanced in tg/wt;*Il10^-/-^* mice compared to fl/fl;*Il10^-/-^* controls and GF mice, quantified as the number of Ki67-positive cells per crypt (Fig. 4H and supplemental Fig. 4D). To validate the loss of mucin-filled GCs as a key characteristic in our CAC mouse model, mucin-filled GCs in the tumor susceptible colon region were assessed as before. The number of mucin-filled GCs was reduced in colonized tg/wt;*Il10^-/-^* mice compared to colonized fl/fl;*Il10^-/-^* and GF mice, with selected crypts in tumor regions again showing a complete loss of mucin-filled GCs (Fig. 4I and supplemental Fig. 4D).

**Fig. 4.**
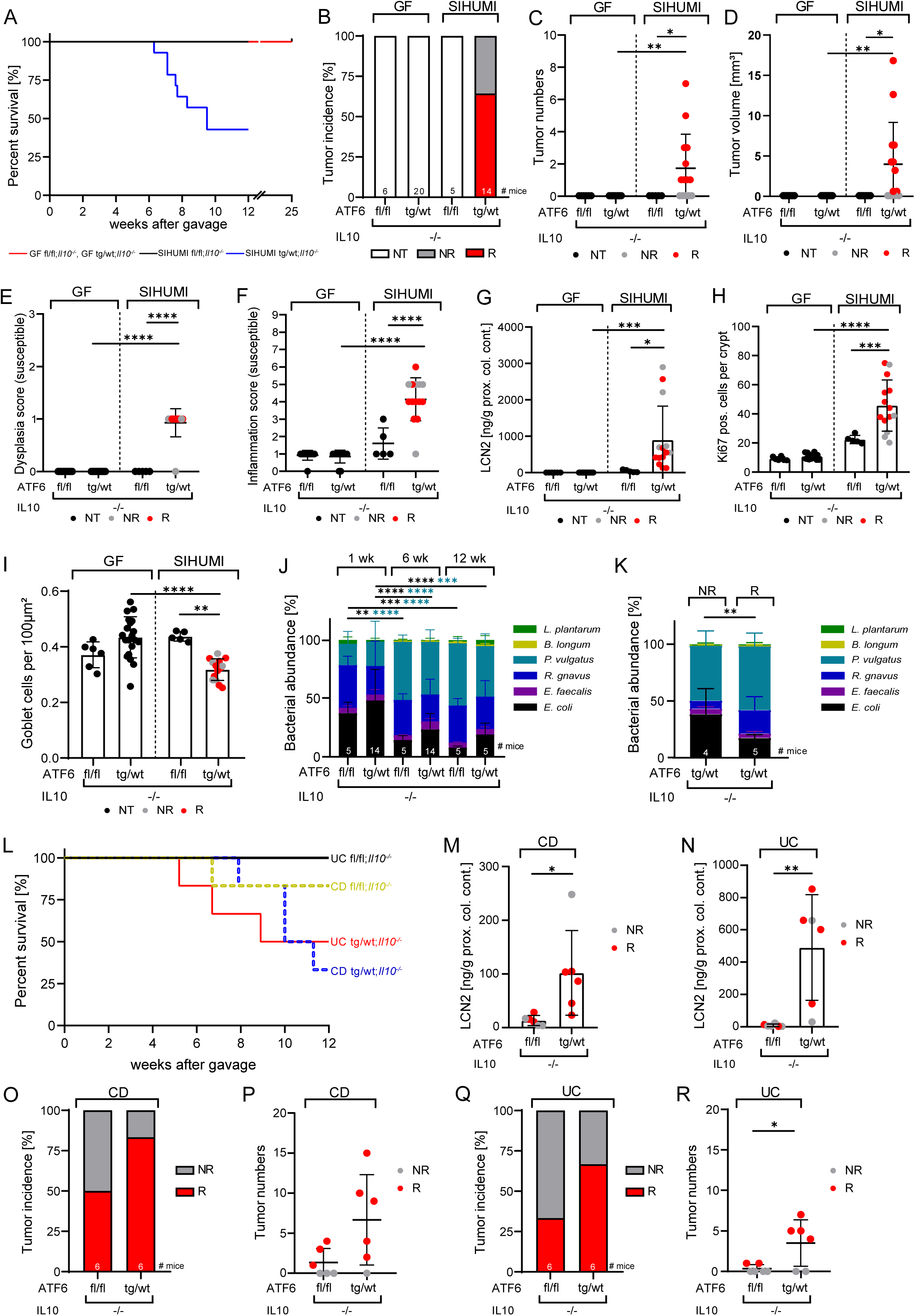
Colonization with an IBD-relevant minimal consortium induces tumor formation in germ-free mice. **A)** Survival curve of germ-free (GF) and SIHUMI colonized nATF6^IEC^;*Il10^-/-^* mice. GF mice were sampled at 25 weeks of age. For SIHUMI colonization, mice were gavaged at the age of four weeks and colonized for 12 weeks. **B)** Colonic tumor incidence of nATF6^IEC^;*Il10^-/-^* mice categorized into non-responder (NR), responder (R) or non-tumor (NT). **C)** Colonic tumor numbers and **D)** volume in nATF6^IEC^;*Il10^-/-^* mice at endpoint. For tumor volume, only the biggest tumor was considered. **E)** Dysplasia score of susceptible colon tissue, resulting from histologically scoring of GF and SIHUMI colonized nATF6^IEC^;*Il10^-/-^* mice for mucosal architecture and atypia (0-3), and **F)** inflammation score (0-9), based on immune cell infiltration and epithelial damage. **G)** Lipocalin-2 (LCN2) levels in proximal colonic content of GF and SIHUMI colonized nATF6^IEC^;*Il10^-/-^* mice. Quantifications of **H)** proliferating cells (Ki67) per crypt and **I)** mucin-filled goblet cells (GC) per 100 µm^2^ in susceptible colon tissue of GF and SIHUMI colonized mice. **J)** Bacterial abundance of SIHUMI strains in mice at week 1, 6 and 12 after gavage. **K)** Bacterial abundance of SIHUMI strains colonized in tg/wt;*Il10^-/-^* separated according to R and NR after 8 weeks of gavage (the time point where most mice dropped out of the experiment). **L)** Survival curve of humanized nATF6^IEC^;*Il10^-/-^* mice (one CD patient, one UC patient). Mice were gavaged at the age of four weeks and colonized for 12 weeks. LCN2 levels in susceptible colonic content of **M)** CD and **N)** UC colonized mice. **O)** Colonic tumor incidence categorized into NR and R and **P)** tumor numbers of CD colonized mice. **Q)** Colonic tumor incidence categorized into NR and R and **R)** tumor numbers of UC colonized mice. GF = germ-free; SIHUMI = simplified human microbiota; NT = non-tumor; NR = non-responder; R = responder; wk = week; CD = Crohn’s disease; UC= ulcerative colitis.

To specify the contribution of the single SIHUMI strains, individual bacterial abundance was determined via qRT-PCR. One week after colonization, *E. coli* comprised approximately 50% of the SIHUMI composition in both fl/fl;*Il10^-/-^* and tg/wt;*Il10^-/^* mice (Fig. 4J). This abundance decreased significantly over time, correlating with a bloom of *P. vulgatus* from six weeks onwards. In the five tg/wt;*Il10^-/-^* mice that survived until week 12 (experimental endpoint), *Lactobacillus plantarum* and *E. coli* showed a tendency towards being more abundant in tg/wt;*Il10^-/-^* mice compared to fl/fl;*Il10^-/-^* mice. Overall composition was otherwise similar between both genotypes at week 12 (Fig. 4J). Interestingly, R and NR tg/wt;*Il10^-/-^* mice showed similar bacterial compositions at the six week pre-tumor time point (supplemental Fig. 4E), and the experimental endpoint at week 12 (supplemental Fig. 4F). At week eight, the average time point where most of the mice with reduced survival dropped out of the experiment, *E. coli* was significantly less abundant in the R tg/wt;*Il10^-/-^* than in NR tg/wt;*Il10^-/-^* mice (Fig. 4K). Moreover, there was a trend towards increased abundance of the mucin-degrading bacterium *Ruminococcus gnavus* in R tg/wt;*Il10^-/-^* mice. However, *R. gnavus* was completely absent in two of the NR tg/wt;*Il10^-/-^* mice (Fig. 4K).

To more closely mimic the human situation in our CAC mouse model, we lastly colonized nATF6^IEC^;*Il10^-/-^* mice with fecal microbiota from CD and UC patients from the aforementioned cohort. Stool samples from two patients with active disease and colonic involvement, as well as matched medication, were chosen for colonization. Mice colonized with CD patient stool showed a reduced survival in both genotypes, with one fl/fl;*Il10^-/-^* and four tg/wt;*Il10^-/-^* mice reaching abortion criteria beginning at approximately seven weeks post gavage. When colonized with UC stool, only three tg/wt;*Il10^-/-^* mice had a reduced survival rate starting from five weeks post gavage (Fig. 4L). Inflammation was verified on protein level, with LCN2 levels being strongly increased in both R and NR tg/wt;*Il10^-/-^* mice compared to fl/fl;*Il10^-/-^* controls for both patient colonizations (Fig. 4M and 4N). Interestingly, tumor formation was observed not only in tg/wt;*Il10^-/-^* mice, but also in fl/fl;*Il10^-/-^* mice, indicating that changes of the microbial composition of IBD patients can already trigger tumorigenesis in an inflamed environment, independent of activated ATF6 (Fig. 4O and 4Q). However, tumors were clearly smaller and less abundant in humanized fl/fl;*Il10^-/-^* mice compared to tg/wt;*Il10^-/-^* mice. Tumor incidence in CD colonized mice was moderately higher than in UC colonized mice, however UC tg/wt;*Il10^-/-^* mice showed a reduced survival compared to CD tg/wt;*Il10^-/-^* mice. 50 % of CD colonized fl/fl;*Il10^-/-^* and 83.3% of tg/wt;*Il10^-/-^* mice developed tumors in the colon (Fig. 4O). In CD colonized fl/fl;*Il10^-/-^* mice, tumor numbers were relatively low (up to four tumors; Fig. 4P) and small in size (up to 2 mm^3^; supplement Fig. 4G) whereas in CD colonized tg/wt;*Il10^-/-^* mice, up to 15 tumors per mouse developed, reaching a volume of 15 mm^3^ (Fig. 4P and supplemental Fig. 4G). In UC colonized mice, tumor incidences were 33.3% for fl/fl;*Il10^-/-^* and 66.7% for tg/wt;*Il10^-/-^* mice (Fig. 4Q). In UC colonized fl/fl;*Il10^-/-^* mice, two mice had one tumor each, the larger with a volume of 4 mm^3^, whereas tumor numbers in UC colonized tg/wt;*Il10^-/-^* mice were higher (up to seven) and larger still (up to 25 mm^3^; Fig. 4R and supplemental Fig. 4H). In summary, these data clearly show that tumorigenesis in the monoallelic nATF6^IEC^;*Il10^-/-^* model is microbiota-dependent, and that inflammation alone is enough to induce tumorigenesis following IBD patient fecal transfer.

## Discussion

Individuals with IBD are at an increased risk of developing CRC, with a resulting poorer prognosis than those with sporadic CRC.^16,17^ The rising prevalence of IBD suggests CAC may represent an important emerging health issue.^18^ While alterations in both ATF6 and IL-10 have been associated with CAC, their complex interplay and potentially crucial role in tumorigenesis remain unclear.^12,19^ Here, we identify ATF6 activation as a driver of invasive carcinomas in a chronic inflammatory milieu, presenting a clear risk for the development of CAC in IBD patients. Our novel murine model of CAC, combining ATF6 activation and the loss of IL-10, is characterized by reduced survival and enhanced susceptibility to tumor formation in the tumor susceptible proximal colon. We demonstrate the human relevance of this model by showing increased cytoplasmic ATF6 expression in patients with active compared to inactive IBD. Furthermore, in support of its translational relevance we recapitulate disease, following colonization of otherwise tumor-free GF mice with a humanized minimal-microbial consortium or fecal microbiota from either CD or UC patients.

We previously identified a role for biallelic ATF6 activation (tg/tg nATF6^IEC^) in microbiota-dependent intestinal tumor formation without the initial presence of inflammation. In the absence of an additional trigger, monoallelic (tg/wt) mice never developed tumors.^8^ Short-term DSS treatment induced tumors in 80% of tg/wt mice, identifying inflammation as a risk factor in ATF6-driven tumorigenesis.^8^ However, the consequence of sustained inflammation remained unclear. To examine the role of chronic inflammation in this context, we crossed nATF6^IEC^ mice with *Il10^-/-^* mice. Chronic inflammation represents an additional burden, as evidenced by enhanced leukocyte infiltration in tumor-susceptible regions, and increased fecal concentrations of LCN2, that clearly enhanced tumor susceptibility in tg/wt;*Il10^-/-^* mice. However, our analyses show that histological inflammation was not markedly enhanced in tg/wt;*Il10^-/-^* mice compared to fl/fl;*Il10^-/-^* controls, indicating that mechanisms besides inflammation contribute to tumor formation in this model. In addition to its immunosuppressive function, IL-10 plays an important role in intestinal epithelial homeostasis, with roles in maintaining epithelial barrier integrity, IEC proliferation and differentiation, as well mucus production.^20–22^ In SPF tg/wt;*Il10^-/-^* mice, we observed an increase in proliferation, with a concomitant reduction in mucin-filled GCs and decreased distance of the mucosal microbiota to the epithelium. IL-10 is involved in maintaining ER folding capacity upon ER stress and is critical in highly secretory cells such as GCs.^22^ The combination of IL-10 deficiency and unresolved ER stress therefore likely substantially diminishes mucus production capacity. Mucus segregates potentially immunostimulatory microbes from the epithelium, and thus disruption of this barrier, particularly in concert with enhanced bacterial penetration, may precipitate disease.^23^ Abrogation of the main mucus glycoprotein *Muc2* in mice for example, leads to an increased tendency of spontaneous colonic tumor formation.^24^ Enhanced bacterial penetration of the mucus layer increases the bacterial interaction potential with the epithelium, and has been identified in various experimental models of CAC as well as in UC patients.^25–27^ Intriguingly, GF tg/wt;*Il10^-/-^* mice did not display a reduction in mucin-filled GC number, nor an increase in proliferation, confirming the requirement of the intestinal microbiota for this phenotype. Crucially, GF tg/wt;*Il10^-/-^* also remained tumor-free, revealing bacterial presence as necessary for tumorigenesis.

To assess the role of the microbiota in tumor formation in tg/wt;*Il10^-/-^* mice, we profiled luminal and mucosal microbiota across different stages of tumor progression. We identified significant alterations in mucosal microbiota between NR and R tg/wt;*Il10^-/-^* mice that were not present in luminal data. Further supporting this finding, machine learning models trained on mucosal data could better predict phenotype than luminal data, demonstrating that mucosal changes are more relevant for disease. Differential abundance analysis between fl/fl;*Il10^-/-^* and R tg/wt;*Il10^-/-^* mice revealed an increase in putative pathobionts such as Mucispirillum, Bacteroides and Desulfovibrio. Intriguingly, multiple species from these genera have previously been associated with both colitis and tumorigenesis in animal models and patients.^28–32^ Bacteroides spp. strongly correlated with tumor number and additionally correlated with ATF6 expression in patients. *Phocaeicola vulgatus* (formerly *Bacteroides vulgatus*), has been shown to potentiate colitis in UC through protease-induced barrier dysfunction.^30^ Additionally, *P. vulgatus* has been demonstrated to accelerate tumor progression via bile salt hydrolase activity, leading to increased β-catenin activation and recruitment of immunosuppressive Foxp3^+^ T_reg_.^31^ Importantly, *P. vulgatus* is present in the modified SIHUMI consortium and was amongst the dominant strains in both genotypes from week 6 onwards, warranting further research into its potential tumor-driving function and mechanistic effects on immune cells.

To our knowledge, spontaneous microbiota-dependent models of CAC are currently sparse,^27,33^ underscoring the pressing need for appropriate animal models to elucidate microbe-host interactions in the context of CAC. We developed and characterized a novel, human-relevant, microbiota-driven model of spontaneous CAC, which will pave the way for future investigations of host-microbe interactions involved in CAC pathogenesis. Such microbiota-dependent models are indispensable for clinically translating microbial involvement in CRC and CAC and enable dissection of host and microbiota-related mechanisms in complex systems.

## Acknowledgements

The Technical University of Munich provided support through the Core Facility Microbiome of the ZIEL Institute for Food & Health for 16S rRNA gene amplicon sequencing and through the Core Facility Comparative Experimental Pathology of the School of Medicine and Health. Furthermore, we thank Christian Jobin for providing the *E. coli* NC101 strain.

## Author contributions

Conceptualization, D.H.; methodology, J.K., A.S., S.K., V.K., M.A., S.J., E.M.R and O.I.C; formal analysis, J.K., A.S., S.K., V.K., S.J., M.R., A.M. and O.I.C; writing – original draft, J.K., A.S., O.I.C. and D.H.; supervision, D.B., M.A., K.S., B.S., O.I.C. and D.H.; funding acquisition D.H.

## Supplemental information titles and legends

**Supplemental Fig. 1.**
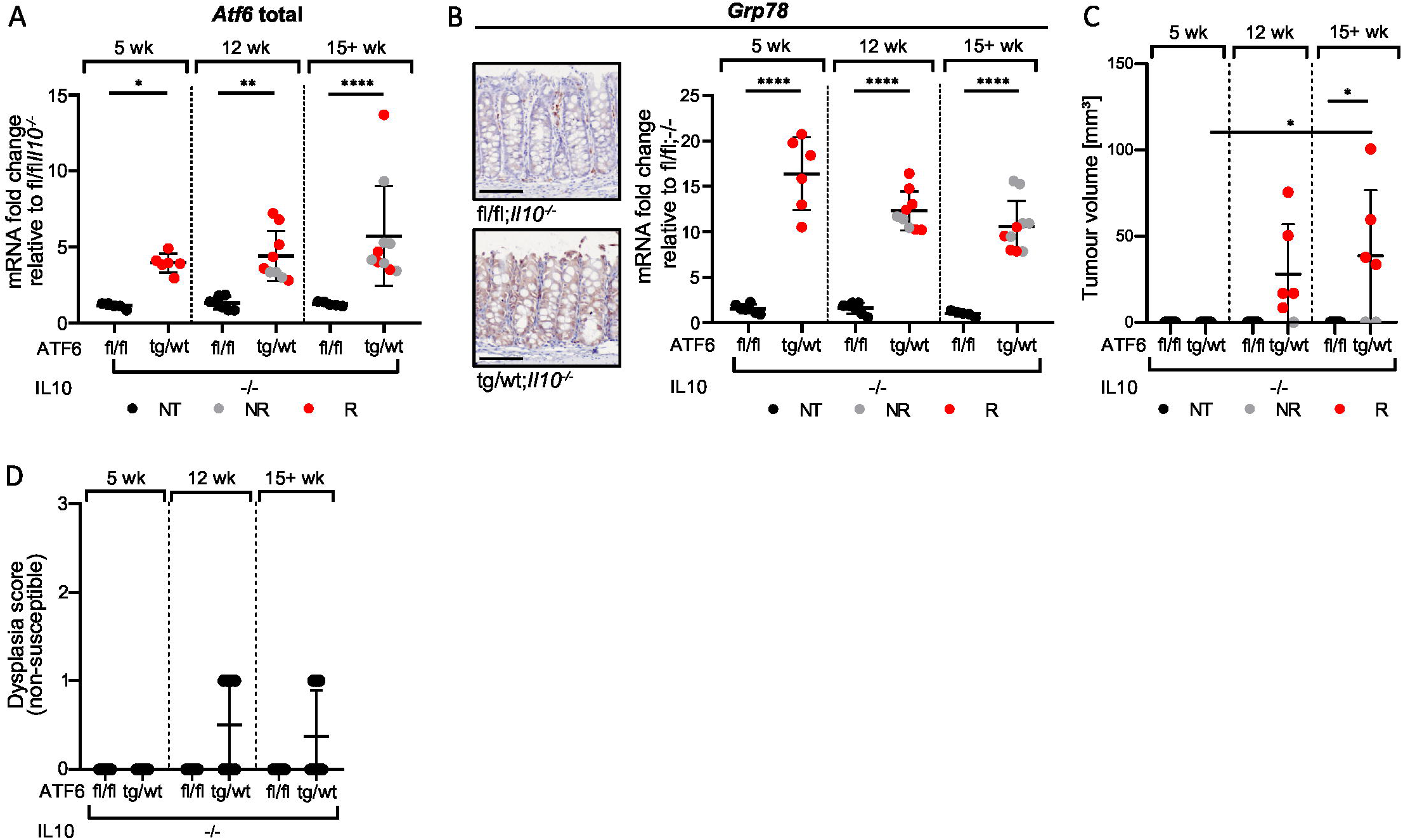
**A)** *Atf6* gene expression levels in whole colon tissue of nATF6^IEC^;*Il10^-/-^* mice at week 5, 12, and 15+. **B)** Representative images (scale bars: 100 µm) and respective gene expression levels of the ER chaperone *Grp78* in whole colon tissue of nATF6^IEC^;*Il10^-/-^* mice at week 5, 12, and 15+. **C)** Colonic tumor volume of nATF6^IEC^;*Il10^-/-^* mice at week 5, 12, and 15+. **D)** Dysplasia score resulting from histologically scoring of non-susceptible colon tissue of nATF6^IEC^;*Il10^-/-^* mice for mucosal architecture and atypia (0-3) at week 5, 12, and 15+. NT = non-tumor; NR = non-responder; R = responder; NUC = nuclear; CYT = cytoplasmic; CD = Crohn’s disease; UC = ulcerative colitis.

**Supplemental Fig. 2.**
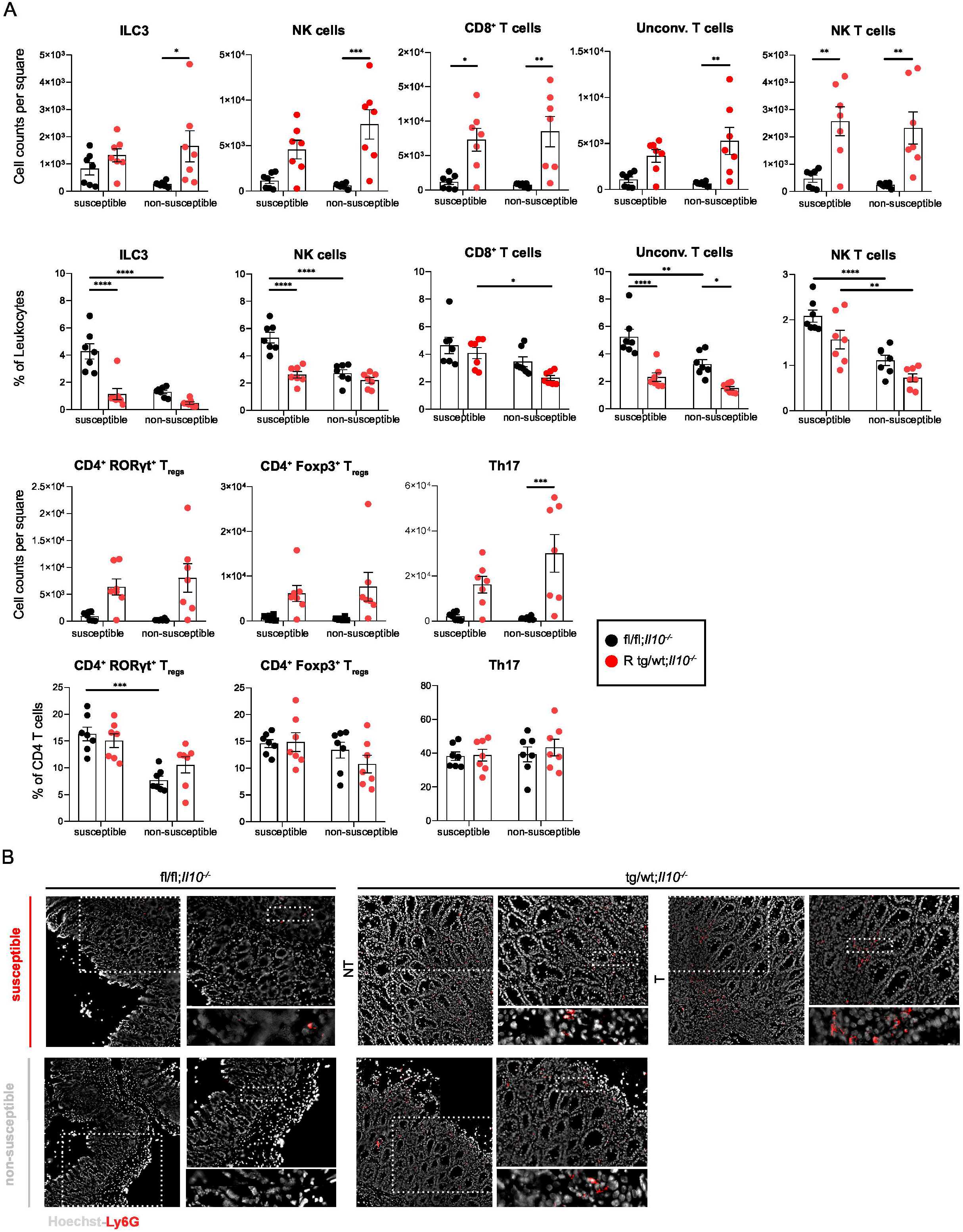
**A)** Quantification of ILC3, NK cells, CD8^+^ cells, unconventional T cells, NK T cells, and different CD4^+^ T_reg_ subpopulations in fl/fl;*Il10^-/-^* control mice and R tg/wt*;Il10^-/-^* mice represented as cell counts per square (0.5 x 0.5 cm colon tissue) and percentage (%) of leukocytes or % of T cells. **B)** Hoechst (grey) and Ly6G (red) staining in the colonic lamina propria of control nATF6^IEC^;*Il10^-/-^* mice and R nATF6^IEC^;*Il10^-/-^* mice under SPF conditions.

**Supplemental Fig. 3.**
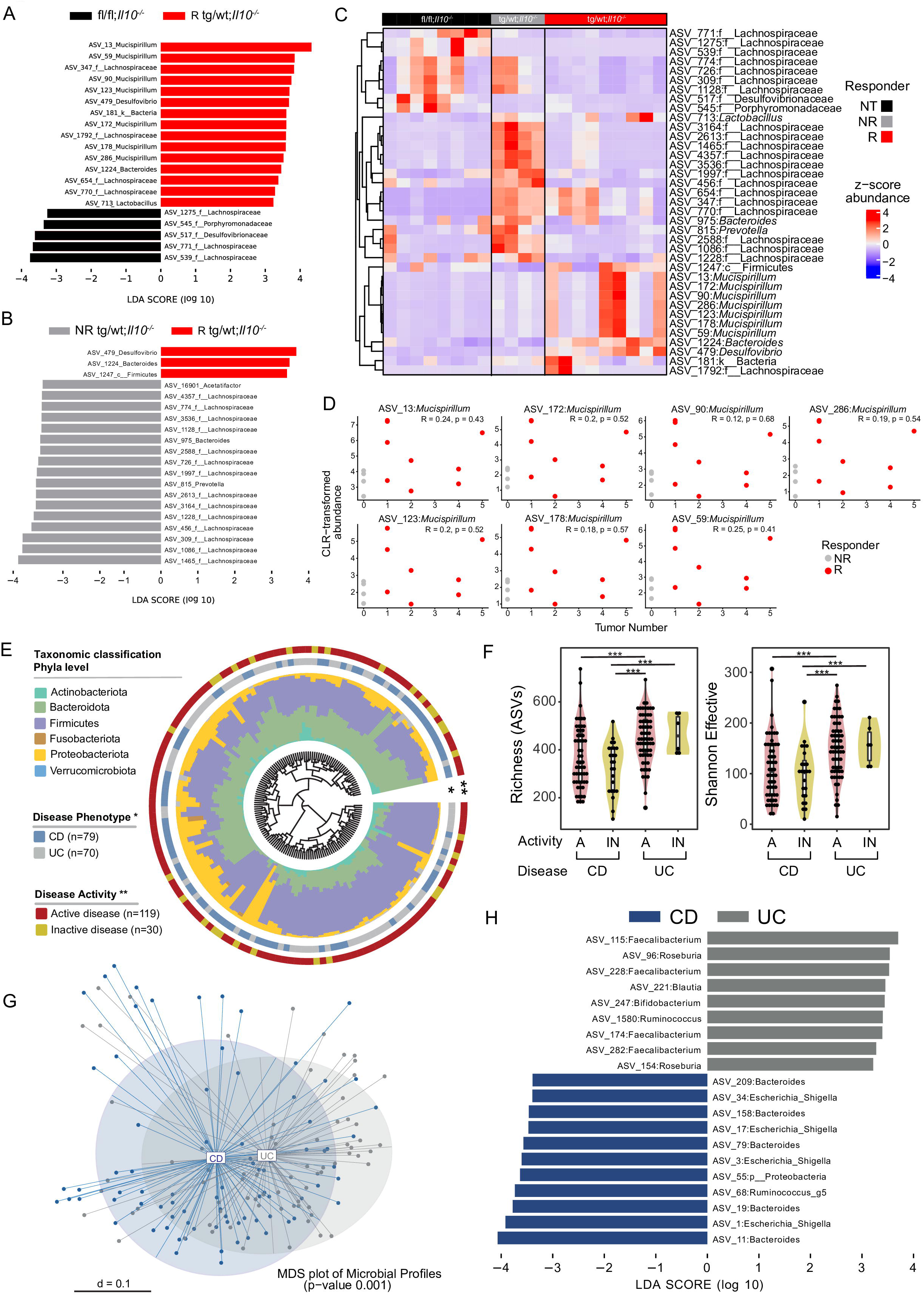
**A)** Differentially abundant taxa between 12-week-old fl/fl;*Il10^-/-^* and R tg/wt;*Il10^-/-^* mice at tumor time points, as identified by LEfSe analysis (LDA threshold = 3.0). **B)** Differentially abundant taxa between 12-week-old NR and R tg/wt;*Il10^-/-^* mice at tumor time points, as identified by LEfSe analysis (LDA threshold = 3.0). **C)** Heatmap depicting z-score of Log_10_ transformed abundances of differentially abundant taxa between fl/fl;*Il10^-/-^* and tg/wt;*Il10^-/-^* NR and R mice. **D)** Spearman-correlation of centered-log-ratio transformed abundance of ASVs classified as Mucispirillum and tumor number in 12-week-old tg/wt;*Il10^-/-^* mice. **E)** Phylogenetic tree showing the similarities between microbiota profiles based on generalized UniFrac distances in mucosal biopsies derived from CD patients (n=79) and UC patients (n=70). Individual taxonomic composition at the phylum level is shown as stacked bar plots around the phylogram. Innermost ring shows stratification based on disease phenotype, CD patient samples (blue) and UC patient samples (grey) and denoted by an asterisk (*); the second ring shows stratification based on disease activity, active disease (red), inactive disease (yellow) and denoted by two asterisks (**). **F)** Alpha-diversity in terms of community richness and effective number of species in mucosa-associated bacteria from CD (blue) and UC (grey) patients, stratified into active (A) and inactive (IN) CD and UC patients. **G)** β-diversity between CD and UC patients based on generalized UniFrac distance. Differences between groups were tested using PERMANOVA. **H)** Differentially abundant taxa between CD and UC patients as identified by LEfSe analysis (LDA threshold = 3.0). NT = non-tumor; NR = non-responder; R = Responder; ASV = amplicon sequence variant.

**Supplemental Fig 4.**
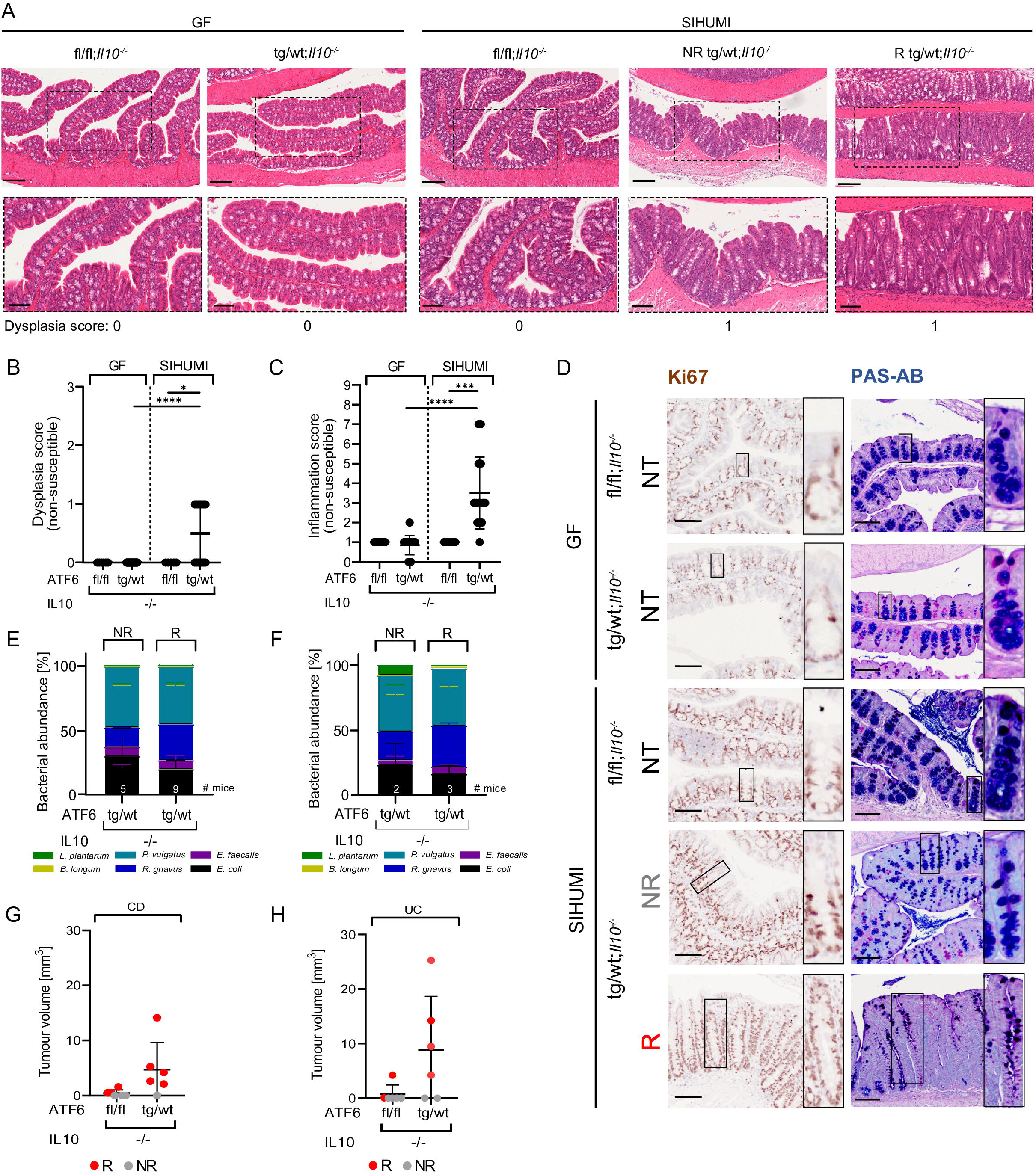
**A)** Representative images of H&E staining of susceptible colon tissue (scale bars: 200 µm) and corresponding higher magnifications (*rectangles*; scale bars: 100 µm) of germ-free (GF) and SIHUMI colonized nATF6^IEC^;*Il10^-/-^* mice. Numbers represent the respective dysplasia score. **B)** Dysplasia score of non-susceptible colon tissue, resulting from histologically scoring of GF and SIHUMI colonized nATF6^IEC^;*Il10^-/-^* mice for mucosal architecture and atypia (0-3), and **C)** inflammation score (0-9), based on immune cell infiltration and epithelial damage. **D)** Ki67 and PAS-AB staining of proximal or susceptible colon tissue of GF and SIHUMI colonized fl/fl;*Il10^-/-^* and tg/wt;*Il10^-/-^* mice (scale bars: 100 µm) and corresponding magnifications (scale bar: 800 µm). Bacterial abundance of SIHUMI strains colonized in tg/wt;*Il10^-/-^* separated according to responder (R) and non-responder (NR) at **E)** week 6 versus **F)** week 12 after gavage. **G)** Colonic tumor volume of CD colonized nATF6^IEC^;*Il10^-/-^* mice. **H)** Colonic tumor volume of UC colonized nATF6^IEC^;*Il10^-/-^* mice. NT = non-tumor; NR = non-responder; R = responder; GF = germ-free; SIHUMI = Simplified human microbiota.

## Materials and Methods

### Ethics statement

All animal experiments were approved by the Animal Health Care and Use Committee of Upper Bavaria (AZ ROB-55.2-1-54-2532-217-14, AZ ROB-55.2-2532.Vet_02-20-58, AZ ROB-55.2-2532.Vet_02-18-121, and AZ ROB-55.2-2532.Vet_02-18-149), and performed in strict compliance with the EEC recommendations for the care and use of laboratory animals (European Communities Council Directive of November 24, 1986 (86/609/EEC)).

nATF6^IEC^ mice were generated as described previously^8^ and subsequently crossed with Interleukin-10 deficient mice (nATF6^IEC^;*Il10^-/-^*). nATF6^IEC^;*Il10^-/-^* mice were housed under SPF and GF conditions (12 h light/dark cycles and 24-26°C) at the Technical University of Munich (Weihenstephan, Germany). Mice were cage-separated by sex but not by genotype, and the breeding scheme enabled generation of all genotypes. All animals received a standard chow diet (Ssniff) *ad libitum*. The standard group size for each experiment was six mice per genotype, unless stated otherwise, and both sexes were used for experiments.

### Colonization of GF mice

For the gavage with a modified version of the Simplified Human Microbiota (SIHUMI; an IBD-related minimal consortium consisting of the five human derived bacteria *Enterococcus faecalis* OG1RF, *Ruminococcus gnavus* ATCC 29149, *Phocaeciola vulgatus* ATCC 8482, *Lactobacillus plantarum* WCFS1, and *Bifidobacterium longum* subsp. *longum* ATCC 15707 and one murine derived *Escherichia coli* NC101), bacteria were grown as described previously.^34^ Briefly, all strains were grown individually in Wilkins-Chalgren anaerobic broth (WCA, Thermo Fisher Scientific) supplemented with 0.05 % (w/v) L-cystein (Carl Roth) and 0.002 % (w/v) dithiothreitol (Sigma Aldrich) in Hungate tubes for 24 h under anaerobic conditions. To ensure equal amounts of each bacterial strain, cells were counted under the microscope and a bacterial mixture was prepared containing approximately 2 x 10^9^ cells/ml. Aliquots were mixed 1:1 with 40 % (v/v) glycerol in WCA medium (previously gassed with N_2_) and sealed with a rubber septum. The bacterial mixture was stored at -80°C until use. The *E. coli* NC101 strain of SIHUMI was kindly provided by Christian Jobin (Department of Medicine, University of Florida, USA).^4^ For the gavage with human stool samples, a 1 ml-aliquot of human stool was thawed on ice and mixed thoroughly before centrifugation for 3 min at 300 x g to remove cell debris. Supernatant was transferred into a new Eppendorf tube and centrifuged for 10 min at 8000 x g at 4°C. The cell pellet was resuspended in 1 ml of reduced PBS (PBS + 0.05% L-Cystein sterile filtered and imported into anaerobic chamber the day before gavage). Gavage liquid was mixed and transferred into sterile Hungate tubes (previously gassed with N_2_), sealed with a rubber septum and used directly for gavage. Preparation of gavage for humanization was performed in an anaerobic chamber (Whitley H85 workstation). For the colonization with SIHUMI or IBD stool samples, GF mice were gavaged once orally at the age of four weeks and colonized for 12 weeks. All mice were euthanized with CO_2_ at the end of an experiment or when abortion criteria were met.

### Tissue processing and sampling of mucosa-associated microbiota

Mice were euthanized, the colon removed immediately and opened longitudinally. Colonic content was removed, tumor number and tumor volume (of the biggest tumor) were measured and the colon processed to a Swiss roll by rolling the tissue from distal to proximal end. Swiss rolls were fixed in 10 % (v/v) formalin for 48 h, dehydrated (Leica TP1020) and embedded in paraffin (McCormick, Leica EG1150C) for histologic analysis. For fluorescence *in situ* hybridization (FISH) analysis, the colon was removed and rolled unopened, before fixation in Carnoy solution (60 % v/v MeOH, 30 % v/v chloroform, 10 % v/v acetic acid) overnight. For dehydration, tissue samples were washed twice with anhydrous MeOH for 30 min, followed by 100 % EtOH for 15 min, 1:1 xylene/100 % EtOH for 5 min and twice xylene for 7 min. Tissue samples were then placed into molten paraffin for 30 min and embedded. For the analysis of the mucosa-associated microbiota, the murine colon was opened longitudinally and colonic content removed. Colon tissue was then washed with sterile PBS to remove residual content. Tumor, tumor-adjacent and non-tumor parts were excised with a sterile scalpel treated with DNA Away^TM^ (Fisher Scientific).

### Histological staining and histopathologic analysis

Formalin-fixed paraffin-embedded (FFPE) colon Swiss rolls were cut into 4-5 µm thick sections and stained with Hematoxylin (of Mayer) and 0.2 % Eosin (H&E, both Medite) with a Leica staining machine (Leica). Colon sections were examined in a blinded manner, as recently described.^35^ Briefly, histological scores for inflammation (based on immune cell infiltration and epithelial damage) and dysplasia (based on degree of mucosal atypia) were assigned, resulting in an inflammation score ranging from 0 (non-inflamed) to 9 (highly inflamed) and a dysplasia score from 0 (no change) to 3 (high grade dysplasia/invasive carcinoma).

For immunohistochemical staining with anti-Ki67 and anti-Grp78, FFPE colon Swiss rolls were cut into 4-5 µm thick sections, deparaffinized and rehydrated. After heat-mediated antigen-retrieval with citrate buffer (10 mM, pH 6) peroxidase quenching was performed (3 % H_2_O_2_, 10 min, Sigma-Aldrich) followed by blocking of the sections with buffer containing 5 % serum of the species in which the secondary antibody was produced. Primary antibody (Anti-Ki67 or Anti-Grp78, both rabbit, 1:300; Cell Signaling Technology) was incubated overnight at 4°C. After three washing steps with PBS, secondary antibody was applied to the sections (1:300, horseradish peroxidase coupled) and incubated for 1 h at RT. Subsequently, antigen detection was performed by DAB/Metal concentrate (Fisher) and nuclei were counterstained with hematoxylin. The PreciPoint M8 microscope (PreciPoint) was used for scanning and analysis. For quantification of anti-Ki-67 staining, the number of Ki-67 positive cells were counted in 20 crypts of the proximal colon.

### Lipocalin-2 ELISA

To generate the supernatant of intestinal content (either from feces or proximal colonic content) for ELISA measurements, a small portion of frozen samples were removed using a sterile spatula and diluted 1:10 in sterile PBS (based on weight), vortexed and incubated in the fridge overnight. On the next day, samples were centrifuged for 5 min at 400 x g to pellet cell debris. The supernatant was transferred into a new Eppendorf tube before centrifuging again for 10 min at 5000 x g to pellet bacteria. The resulting supernatant was again transferred into a new Eppendorf tube and stored at -80°C until used. Lipocalin-2 was measured using the DuoSet^TM^ ELISA development system (R&D systems) according to manufacturer’s instructions. Concentrations were determined using the standard curve method and expressed as nanograms per gram of intestinal content.

### Immune cell isolation and flow cytometry

Murine colon was isolated, all fat was removed and the colon was flushed with Gut wash solution (HBSS w/o Ca/Mg (Sigma-Aldrich) 5 mM EDTA (Fisher Bioreagents), 5 % FCS (Sigma), 10 mM HEPES (Sigma-Aldrich)) before being cut open longitudinally. Proximal (2/3) and distal (1/3) parts were separated, and feces and mucus removed. Each part was cut into pieces, placed in Gut wash solution containing 1 mM DTT (Sigma) and incubated for 30 min at 37°C while shaking at 170 rpm. After vortexing for 10 sec, tissue was transferred into fresh Gut wash containing 1 mM DTT and incubated for as before. Following this, colon pieces were washed with Gut digestion solution (HBSS with Ca/Mg (Sigma-Aldrich), 5 % FCS (Sigma), 10 mM HEPES) for 15 min at 4°C. Remaining tissue was minced and digested in Gut digestion solution supplemented with 400 U/ml CollagenaseIV (Worthington) and 0.4 mg/ml DNaseI (Roche) for 30 min at 37°C while shaking at 125 rpm. Cells were meshed through a 100 µm strainer and washed once with Gut medium (RPMI (Sigma-Aldrich), 10 % FCS (Sigma), 1 % antibiotic-antimycotic solution (100x; including 10000 units penicillin, 10 mg streptomycin and 25 µg amphotericin B per ml; Sigma), 1% L-Glutamine (Sigma), 10 mM HEPES, 0.05 mM β-Mercaptoethanol (Gibco)). Leukocytes were enriched over a 70 % - 37 % - 30 % gradient, prepared from isotonic percoll solution (90 % Percoll (Cytiva), 10 % 10x PBS (Gibco). The interphase between 70 % and 37 % was collected, washed once with Gut medium, and resuspended in FACS buffer (PBS (Sigma), 1 % FCS (Sigma), 2.5 mM EDTA (Invitrogen), 0.02 % sodium azide (Sigma)).

For Live/Dead staining, cells were washed twice with PBS, resuspended in 100 µl PBS containing Zombie aqua (1:1000; BioLegend), and incubated for 15 min at 4°C. Cells were washed twice, followed by Fc Block (anti-mouse purified CD16/32 in FACS buffer for 10 min at 4°C. Surface staining was performed in 100 µl for 20 min at 4°C. Cells were washed twice using FACS buffer, followed by fixation and intranuclear staining using ebioscience Foxp3 transcription factor staining kit (Invitrogen).

Data was collected on a Cytek® Aurora, and datasets were further analyzed in FlowJo^TM^ 10 (Tree Stra, Inc). CountBrightTM Absolute counting Beads (Thermo Fisher Scientific) were used for total cell quantification. Results are shown in cell counts per square (0.5 cm^2^) and % leukocytes.

Gating was performed as follows: leukocytes were pre-gated as CD45.2^+^ live, singlets, followed by separation into a CD90.2^+^ fraction for T cells and ILCs and a CD90.2^-^ fraction for myeloid cells. Within the CD90.2^-^ fraction, neutrophils were identified as CD11b^+^Ly6G^+^ and from Ly6G^-^ cells, monocytes were gated as CD11b^high^ and Ly6C^+^. From Ly6C^-^ fraction CD11c high to low, MHCII^+^ cells were selected, and CD64 expressing macrophages separated from CD64^-^CD11c^high^ conventional dendritic cells (cDCs). The latter were further subdivided into CD103^+^ cDC1, CD11b^+^ cDC2, and intestine-characteristic CD103^+^CD11b^+^ cDC2. Furthermore, a small fraction of CD103^-^ CD11b^-^DCs were identified. Within CD90.2^+^ cells, NK cells were gated as NK1.1^+^, being CD3ε^-^. CD3ε^+^ NK1.1 cells were identified as NK T cells. From NK1.1^-^ cells, T cells were identified through their CD3ε and CD90.2 expression and subdivided by CD4 and CD8a into CD4^+^ T cells, CD8^+^ T cells, and unconventional T cells lacking CD4 and CD8a expression. Regulatory T cells were identified within the CD4^+^ T cells through their expression of Foxp3 and further subdivided into gut-specific Rorgt+ Treg and classical Tregs expressing only Foxp3. The remaining CD90.2^+^ CD3ε^-^ cells were identified as ILCs; hereby, Rorgt^+^ ILCs were identified as ILC3 and Tbet^+^ ILCs as ILC1s.

### ChipCytometry

ChipCytometry staining was generated as described by Jarosch *et al*.^36^ Briefly, cryo-preserved colon Swiss rolls were cut into 7 µm thick sections and placed on a coverslip. Samples were fixed for 45 min with fixation buffer (Zellkraftwerk Chip Kit) and washed before transferring the coverslip to the chip. Sections were scanned and each position was photo-bleached for 20 sec using the ChipCytometry microscope (ZellscannerONE/Zeiss Axio Observer 7) to reduce the auto-fluorescence of the sample. Next, a background image was acquired in all channels using the ChipCytometry microscope. For the staining, 300 µl antibody master mix per chip (Table 4) was prepared. The antibody master mix was centrifuged for 10 min at 4°C, 16 000 × *g* to remove dye or antibody aggregates. After acquisition of the background image, the chip was removed from the microscope and washed with sterile PBS before the antibody master mix was applied and incubated for 30 min in the dark at RT. Afterwards, the chip was washed with sterile PBS for several rounds (rinsed five times with PBS, washed once with 20 ml PBS and rinsed five times with PBS again), the staining was scanned using the ChipCytometry microscope. To continue with the next panel of staining, each position was again photo-bleached for 20 sec and the background was re-adjusted. After the completion of all stainings, images were adjusted by finding the ideal combination between the contrast and the background in the ZellExplorer App of the instrument. Signal intensity was analyzed as described before.^37^ Quantification was performed by gating the cell populations using flow cytometry software (FlowJo).

**Table 3:** Antibody list for flow cytometry.

| Antigen | Clone | Fluorochrome | Company |
| --- | --- | --- | --- |
| CD3ε | 145-2C11 | APC-Cy7 | BioLegend |
| CD4 | GK1.5 | BUV737 | BioLegend |
| CD8a | 53-6.7 | PerCP | BioLegend |
| CD11b | M1/70 | BV570 | BioLegend |
| CD11c | N418 | PB | BioLegend |
| CD16/CD32 | 2.4G2 | purified | BD Biosciences |
| CD44 | IM7 | BV650 | BioLegend |
| CD45.2 | 104 | PE/Dazzle594, PB | BioLegend |
| CD62L | MEL-14 | PE/Cy7 | Tonbo Bioscience |
| CD64 (FcγRI) | X54-5/7.1 | APC | BioLegend |
| CD86 | GL-1 | BV605 | BioLegend |
| CD90.2 | 30-H12 | SB780 | BioLegend |
| CD103 | M290 | BUV395 | BD Biosciences |
| CD161c/NK-1.1 | PK136 | PE/Cy5 | BioLegend |
| CD172a (SIRPα) | P84 | PerCP eFluor710 | BioLegend |
| CD274 (PD-L1) | MIH5 | BUV737 | eBioscience |
| CD279 (PD-1) | J43 | BUV805 | eBioscience |
| I-A/I-E (MHCII) | M5/114.15.2 | AF700 | BioLegend |
| Ly-6C | HK1.5 | FITC | BioLegend |
| LY-6G | 1A8 | PerCP/Cy5.5 | BioLegend |
| FOXP3 | 150D | AF647 | BioLegend |
| ROR gamma (t) | Q31-378 | BV421 | BD Biosciences |
| T-bet | 4B10 | PE | BioLegend |
| Mouse IgG1, κ | MOPC-21 | PE, AF647 | BioLegend |

**Table 4:** Antibody list for ChipCytometry.

| Antibody | Fluorophore | Concentration | Company |
| --- | --- | --- | --- |
| CD3 | PE | 1:200 | BioLegend |
| CD11b | PecP/Cy5.5 | 1:50 | BioLegend |
| CD11c | FITC | 1:100 | Life Technologies |
| Hoechst | BUV395 | 1:100000 | Thermo Fisher Scientific |
| Ly6G | PE | 1:100 | BD |

### Periodic Acid-Schiff Alcian Blue (PAS-AB) staining and FISH

For PAS-AB staining, FFPE colon Swiss rolls were cut into 4-5 µm thick sections, deparaffinized and rehydrated. Acidic mucins were stained with Alcian blue (0.5 % w/v in 3 % v/v acetic acid, pH 2.5; Fisher) for 5 min. After treatment with periodic acid solution (0.5 % v/v, 10 min), co-staining with Schiff’s reagent (Sigma-Aldrich) for 10 min was performed to visualize neutral mucins. Nuclei were counterstained with hematoxylin. For quantification of PAS-AB positive goblet cells, 50 crypts were counted in the proximal colon and displayed as number of goblet cells per 100 µm^2^. For FISH, Carnoy-fixed paraffin-embedded colon snails were cut into 9 µm thick sections, deparaffinized, rehydrated and fixed in 10 % (v/v) formalin. Tissue was permeabilized in lysis buffer (1.2 % v/v Triton-X-100 buffer, 20 mM Tris, 2 mM EDTA supplemented with 40 mg/ml (w/v) lysozyme, Sigma-Aldrich) for 45 min at 37°C. Subsequently, tissue sections were incubated with Cy5-conjugated EUB338 (5’-GCTGCCTCCCGTAGGAGT-3’) or scrambled FITC-labelled nonEUB (5‘-ACATCCTACGGGAGGC-3’) in 100 ml sterile hybridization buffer (20 mM Tris, 0.9 M NaCl, 0.01 % v/v SDS-solution (AppliChem GmbH), pH 7.3) overnight at 46°C. Sections were co-stained with anti-Ulex Europeaus Agglutinin 1 (UEA1, Rhodamine-conjugated, 1:1000; Novus Biologicals) and counterstained with 4,6-diamidino-2-phenylindole (DAPI, 1:1000; Sigma-Aldrich). Bacterial distance to the epithelium was measured at five separate random points within five distinct regions of the proximal colon using the Volocity software (Quorum technologies) and averaged.

### Gene expression analysis

RNA from whole colon Swiss roll sections stored in Optimal cutting temperature (OCT) compound was isolated with the NucleoSpin RNAII Kit (Macherey-Nagel) according to manufacturer’s instructions. Afterwards, complementary DNA was synthesized using the Moloney Murine Leukemia Virus (MMLV) reverse transcriptase point mutant synthesis system (Promega) with random hexamers. For quantification, quantitative real-time PCR (qRT-PCR) was performed using the Light Cycler 480 Universal Probe Library System (Roche Diagnostics). Gene expression levels of *Grp78* and *Atf6 total* (transgene and endogenous *Atf6*) were calculated by the 2^-ΔΔCt^ method and normalized to the housekeeper glyceraldehyde-3-phosphate dehydrogenase (*Gapdh*). Primer sequences and respective UPL probes used are shown in Table 5.

**Table 5:**
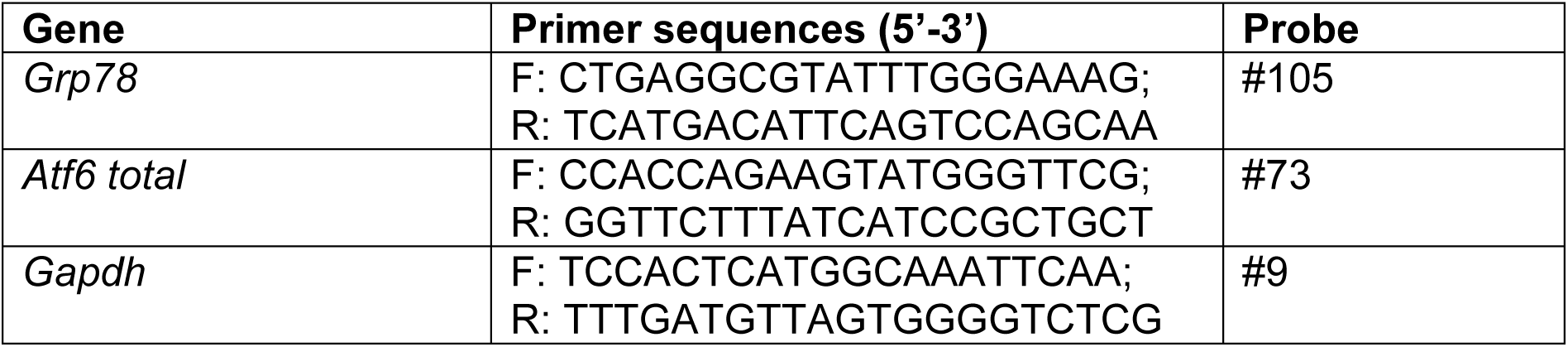
Primer sequences and UPL probes used for quantitative real-time PCR.

### gDNA isolation and 16S rRNA sequencing

DNA isolation from frozen cecal or fecal content was performed as previously described^38^ with minor changes. Briefly, intestinal content was transferred into tubes containing 500 mg silica beads (Lysing Matrix D tubes, MP Biomedicals) and mixed with 250 µl 4 M guanidine thiocyanate, 500 µl 5 % *N*-lauroly sarcosine and 600 µl Stool DNA stabilizer (Invitek Molecular GmbH). Samples were incubated at 70°C for 1 h under constant shaking (700 rpm). Mechanical disruption was achieved using the FastPrep®-24 bead beater (3 x 40 sec, 6.5 m/s; MP Biomedicals). In order to remove phenol contaminations, 15 mg of polyvinylpolypyrrolidone (PVPP; Sigma-Aldrich) was added to the homogenate, vortexed, and centrifuged for 3 min at 15000 x g and 4°C. The supernatant was transferred into a new tube and centrifuged again to acquire a clear supernatant containing lysed bacterial cells. In order to remove bacterial RNA, RNAse (10 mg/ml) was added to 500 µl supernatant and incubated at 37°C for 30 min under constant shaking. The resulting genomic DNA was purified using the NucleoSpin® gDNA clean-up Kit (Macherey-Nagel) according to manufacturer’s instructions. Finally, concentration and purity of the extracted DNA were determined with the Implen NanoPhotometer (Implen GmbH).

### Bacterial community analyses of the minimal consortium SIHUMI

To identify bacterial abundances of the individual strains of the SIHUMI consortium, primers targeting species-specific regions of the 16S rRNA gene were used as previously described.^34^ Briefly, qRT-PCR was performed using the Light Cycler 480 Universal Probe Library System (Roche Diagnostics) and standard curves for each bacterial strain were generated to identify 16S rRNA gene copy numbers per species and per ng DNA in fecal pellets. Bacterial abundance of each strain was then calculated with the formula [(copies of 16S rRNA gene of a specific bacterium/cumulative copies of 16S rRNA gene of all SIHUMI bacteria) x 100] as previously described.^15^

### IBD patient cohort

The ELYP cohort was approved by the French Ethic Committee – Hôptial Saint-Louis (CPP 2016-01-05 RBM) and declared to ClinicalTrials.gov (registration number: NCT02693340). This study was conducted by Prof. Matthieu Allez at the Hôpital Saint-Louis in Paris and written informed consent was obtained from all subjects. In this longitudinal study, 36 Crohn’s disease (CD) and 32 Ulcerative colitis (UC) patients were followed for one year after initiation of biological therapy – either anti-tumor necrosis factor (Infliximab, Golimumab, Adalimumab) or Ustekinumab, Vedolizumab therapy. Clinical disease monitoring and sampling were performed at three time points (week 0, 14, and 52 after initiation of therapy). All patients aged 18 or older with active disease at baseline were included. Samples from patients who did not receive antibiotics at least three months before initiation of biological therapy are included in the analysis.

### Bacterial genomic DNA extraction from IBD patient tissue

Bacterial genomic DNA was extracted using the NucleoSpin® Tissue Kit (Macherey-Nagel). Tissue biopsies were resuspended in 180 µl of sterile-filtered lysis buffer (2 mM EDTA, 1 % Triton X-100 and 20 mM Tris/HCl, pH 8). Freshly prepared lysozyme (20 mg/ml) was added to the suspension, followed by incubation in a shaker at 37°C with 950 rpm for 30 min. Subsequently, Proteinase K (10 mg/ml) was added to the mixture, which was then vortexed vigorously and incubated for 1-3 h at 56°C with 950 rpm until complete tissue lysis. The DNA purification process was completed following the NucleoSpin® Tissue Kit manufacturer’s recommendations.

### High-throughput 16S rRNA gene amplicon sequencing

After genomic DNA extraction, bacterial DNA was diluted in PCR-grade water and used as a template for PCR performed in duplicates. The V3/V4 regions of the 16S rRNA genes were amplified (15 x 15 cycles for biopsies) using bacteria-specific primers 341F and 785R,^39^ followed by a two-step procedure to limit amplification bias.^40^ The concentration of PCR products was measured using fluorometry and adjusted to a 2 nM concentration before pooling. Amplicon purification was conducted using the AMPure XP system (Beckman-Coulter). Sequencing of the pooled samples was performed in paired-end mode (2 x 250 bp) using a MiSeq system (Illumina) according to manufacturer’s guidelines, with 25 % (v/v) PhiX standard library included. Positive controls (a mock community, ZymoBIOMICS, No. D6300) and negative controls (PCR control without DNA template and a DNA extraction control of DNA stabilizer) were included to monitor sequencing artifacts.

### IBD patient tissue staining and quantification

For immunohistochemcial staining of ATF6 in human colon biopsies, anti-ATF6 antibody (1:100; Sigma-Aldrich) was used, following antigen retrieval in EDTA buffer (30 min, pH 9). Hematoxylin (Waldeck) was used to counterstain the nuclei. Staining was performed using the Leica Bond RXm (Wetzlar, Germany) and scanned using the Aperio AT2 (Leica Biosystems, Wetzlar, Germany). For quantification of ATF6-staining, QuPath Software was applied as previously described.^41^ Briefly, cells were identified through positive cell detection of the hematoxylin-positive nucleus. ATF6 staining intensities for the nuclear and the cytoplasmic regions were classified according to defined thresholds of the DAB OD max and DAB OD mean, respectively. Furthermore, the object classifier was trained and applied to distinguish between epithelial cells and stromal cells. Exported epithelial histochemical-scores (H-scores, capturing both the intensity and the percentage of stained cells) for the nucleus and the cytoplasm were used for further patient cohort analyses.

### Machine Learning

Supervised classification of mucosal and luminal microbial profiles was performed using the SIAMCAT R package, utilizing ridge-regression, L1-regularized lasso regression and random forest models. Datasets were adjusted for sample size via random subsampling, low prevalent features were removed, and data was centered log-ratio transformed. All models were trained using repeated leave-one-out CV (LOOCV).

### Statistics

Statistical analyses were performed using R or GraphPad Prism (Version 8.0.1). For the comparison of two groups, an unpaired *t*-test was used. Differences in β diversity were assessed using Permutational multivariate analysis of variance (PERMANOVA).^42^ For multiple comparison, one-way analysis of variance (ANOVA) with Tukey’s multiple comparisons test or two-way ANOVA with Tukey’s multiple comparisons test were used. For the comparison of survival curves Log-rank (Mantel-Cox) test was used. For testing differences in tumor incidences, Chi square test with Fisher’s exact test was used. Data are presented as mean ± standard deviation. *P*-values < 0.05 were considered statistically significant (\**p* < 0.05; \*\**p* < 0.01; \*\*\**p* < 0.001; \*\*\*\**p* < 0.0001).

## References

1. Wang R, Li Z, Liu S, Zhang D. Global, regional and national burden of inflammatory bowel disease in 204 countries and territories from 1990 to 2019: A systematic analysis based on the Global Burden of Disease Study 2019. BMJ Open. 2023;13(3):1–14. doi:10.1136/bmjopen-2022-065186

2. Ahmed M, Metwaly A, Haller D. Modeling microbe-host interaction in the pathogenesis of Crohn’s disease. Int J Med Microbiol. 2021;311(3):151489. doi:10.1016/j.ijmm.2021.151489

3. Cougnoux A, Dalmasso G, Martinez R, et al. Bacterial genotoxin colibactin promotes colon tumour growth by inducing a senescence-associated secretory phenotype. Gut. 2014;63(12):1932–1942. doi:10.1136/gutjnl-2013-305257

4. Arthur JC, Perez-Chanona E, Mühlbauer M, et al. Intestinal inflammation targets cancer-inducing activity of the microbiota. Science (80-). 2012;338(6103):120–123. doi:10.1126/science.1224820

5. Metwaly A, Reitmeier S, Haller D. Microbiome risk profiles as biomarkers for inflammatory and metabolic disorders. Nat Rev Gastroenterol Hepatol. 2022;19(6):383–397. doi:10.1038/s41575-022-00581-2

6. Flemer B, Lynch DB, Brown JMR, et al. Tumour-associated and non-tumour-associated microbiota in colorectal cancer. Gut. 2017;66(4):633–643. doi:10.1136/gutjnl-2015-309595

7. Zeller G, Tap J, Voigt AY, et al. Potential of fecal microbiota for early-stage detection of colorectal cancer. Mol Syst Biol. 2014;10(11):1–18. doi:10.15252/msb.20145645

8. Coleman OI, Lobner EM, Bierwirth S, et al. Activated ATF6 Induces Intestinal Dysbiosis and Innate Immune Response to Promote Colorectal Tumorigenesis. Gastroenterology. 2018;155(5):1539–1552.e12. doi:10.1053/j.gastro.2018.07.028

9. Grootjans J, Kaser A, Kaufman RJ, Blumberg RS. The unfolded protein response in immunity and inflammation. Nat Rev Immunol. 2016;16(8):469–484. doi:10.1038/nri.2016.62

10. Wang M, Kaufman RJ. The impact of the endoplasmic reticulum protein-folding environment on cancer development. Nat Rev Cancer. 2014;14(9):581–597. doi:10.1038/nrc3800

11. Stengel ST, Fazio A, Lipinski S, et al. Activating Transcription Factor 6 Mediates Inflammatory Signals in Intestinal Epithelial Cells Upon Endoplasmic Reticulum Stress. Gastroenterology. 2020;159(4):1357–1374.e10. doi:10.1053/j.gastro.2020.06.088

12. Hanaoka M, Ishikawa T, Ishiguro M, et al. Expression of ATF6 as a marker of pre-cancerous atypical change in ulcerative colitis-associated colorectal cancer: a potential role in the management of dysplasia. J Gastroenterol. 2018;53(5):631–641. doi:10.1007/s00535-017-1387-1

13. Chen J, Bittinger K, Charlson ES, et al. Associating microbiome composition with environmental covariates using generalized UniFrac distances. Bioinformatics. 2012;28(16):2106–2113. doi:10.1093/bioinformatics/bts342

14. Urbauer E, Aguanno D, Mindermann N, et al. Mitochondrial perturbation in the intestine causes microbiota-dependent injury and gene signatures discriminative of inflammatory disease. Cell Host Microbe. 2024;32(8):1347–1364.e10. doi:10.1016/j.chom.2024.06.013

15. Eun CS, Mishima Y, Wohlgemuth S, et al. Induction of bacterial antigen-specific colitis by a simplified human microbiota consortium in gnotobiotic interleukin-10-/-mice. Infect Immun. 2014;82(6):2239–2246. doi:10.1128/IAI.01513-13

16. Shah SC, Itzkowitz SH. U . S . Department of Veterans Affairs. 2023;162(3):715–730. doi:10.1053/j.gastro.2021.10.035.Colorectal

17. Birch RJ, Burr N, Subramanian V, et al. Inflammatory Bowel Disease-Associated Colorectal Cancer Epidemiology and Outcomes: An English Population-Based Study. Am J Gastroenterol. 2022;117(11):1858–1870. doi:10.14309/ajg.0000000000001941

18. Lu C, Schardey J, Zhang T, et al. Survival Outcomes and Clinicopathological Features in Inflammatory Bowel Disease-associated Colorectal Cancer: A Systematic Review and Meta-analysis. Ann Surg. 2022;276(5):e319–e330. doi:10.1097/SLA.0000000000005339

19. Kühn R, Löhler J, Rennick D, Rajewsky K, Müller W. Interleukin-10 deficient mice develop chronic enterocolitis. Cell. 1993;75:263–274.

20. Landuyt AE, Klocke BJ, Duck LW, et al. ICOS Ligand and IL-10 Synergize to Promote Host-Microbiota Mutualism. Proc Natl Acad Sci U S A. 2021;118(13):e2018278118. doi:10.1073/pnas.2018278118

21. Jenkins BR. Loss of Interleukin-10 Receptor Disrupts Intestinal Epithelial Cell Proliferation and Skews Differentiation towards the Goblet Cell Fate. FASEB J. 2021;35. doi:10.1096/fj.202002369R

22. Hasnain SZ, Tauro S, Das I, et al. IL-10 promotes production of intestinal mucus by suppressing protein misfolding and endoplasmic reticulum stress in goblet cells. Gastroenterology. 2013;144(2):357–368.e9. doi:10.1053/j.gastro.2012.10.043

23. Hansson GC, Johansson MEV. The inner of the two Muc2 mucin-dependent mucus layers in colon is devoid of bacteria. Gut Microbes. 2010;1(1):51–54. doi:10.4161/gmic.1.1.10470

24. Van der Sluis M, De Koning B a. E, De Bruijnp ACJM, et al. Muc2-Deficient Mice Spontaneously Develop Colitis, Indicating That MUC2 Is Critical for Colonic Protection. Gastroenterology. 2006;131(1):117–129. doi:10.1053/j.gastro.2006.04.020

25. Johansson MEV, Gustafsson JK, Holmen-Larsson J, et al. Bacteria penetrate the normally impenetrable inner colon mucus layer in both murine colitis models and patients with ulcerative colitis. Gut. 2014;63(2):281–291. doi:10.1136/gutjnl-2012-303207

26. van der Post S, Jabbar KS, Birchenough G, et al. Structural Weakening of the Colonic Mucus Barrier Is an Early Event in Ulcerative Colitis Pathogenesis. Gut. 2019;68(12):2142–2151. doi:10.1136/gutjnl-2018-317571

27. Slowicka K, Petta I, Blancke G, et al. Zeb2 Drives Invasive and Microbiota-Dependent Colon Carcinoma. Nat Cancer. 2020;1(6):620–634. doi:10.1038/s43018-020-0070-2

28. Kuffa P, Pickard JM, Campbell A, et al. Fiber-Deficient Diet Inhibits Colitis through the Regulation of the Niche and Metabolism of a Gut Pathobiont. Cell Host \& Microbe. 2023;31(12):2007–2022.e12. doi:10.1016/j.chom.2023.10.016

29. Yazici C, Wolf PG, Kim H, et al. Race-dependent association of sulfidogenic bacteria with colorectal cancer. Gut. 2017;66(11):1983–1994. doi:10.1136/gutjnl-2016-313321

30. Mills RH, Dulai PS, Vázquez-Baeza Y, et al. Multi-Omics Analyses of the Ulcerative Colitis Gut Microbiome Link Bacteroides Vulgatus Proteases with Disease Severity. Nat Microbiol. 2022;7(2):262–276. doi:10.1038/s41564-021-01050-3

31. Sun L, Zhang Y, Cai J, et al. Bile Salt Hydrolase in Non-Enterotoxigenic Bacteroides Potentiates Colorectal Cancer. Nat Commun. 2023;14(1):755. doi:10.1038/s41467-023-36089-9

32. Xie R, Gu Y, Li M, et al. Desulfovibrio Vulgaris Interacts with Novel Gut Epithelial Immune Receptor LRRC19 and Exacerbates Colitis. Microbiome. 2024;12(1):4. doi:10.1186/s40168-023-01722-8

33. Han JX, Tao ZH, Qian Y, et al. ZFP90 drives the initiation of colitis-associated colorectal cancer via a microbiota-dependent strategy. Gut Microbes. 2021;13(1):1–20. doi:10.1080/19490976.2021.1917269

34. Lengfelder I, Sava IG, Hansen JJ, et al. Complex bacterial consortia reprogram the colitogenic activity of enterococcus faecalis in a gnotobiotic mouse model of chronic, immune-mediated colitis. Front Immunol. 2019;10(JUN):1–18. doi:10.3389/fimmu.2019.01420

35. Remke M, Groll T, Metzler T, et al. Histomorphological scoring of murine colitis models : A practical guide for the evaluation of colitis and colitis-associated cancer. Exp Mol Pathol. 2024;140(September). doi:10.1016/j.yexmp.2024.104938

36. Jarosch S, Köhlen J, Wagner S, D’Ippolito E, Busch DH. ChipCytometry for multiplexed detection of protein and mRNA markers on human FFPE tissue samples. STAR Protoc. 2022;3(2). doi:10.1016/j.xpro.2022.101374

37. Jarosch S, Köhlen J, Sarker RSJ, et al. Multiplexed imaging and automated signal quantification in formalin-fixed paraffin-embedded tissues by ChipCytometry. Cell Reports Methods. 2021;1(7). doi:10.1016/j.crmeth.2021.100104

38. Godon JJ, Zumstein E, Dabert P, Habouzit F, Moletta R. Molecular microbial diversity of an anaerobic digestor as determined by small-subunit rDNA sequence analysis. Appl Environ Microbiol. 1997;63(7):2802–2813. doi:10.1128/aem.63.7.2802-2813.1997

39. Klindworth A, Pruesse E, Schweer T, et al. Evaluation of general 16S ribosomal RNA gene PCR primers for classical and next-generation sequencing-based diversity studies. Nucleic Acids Res. 2013;41(1):1–11. doi:10.1093/nar/gks808

40. Berry D, Mahfoudh K Ben, Wagner M, Loy A. Barcoded primers used in multiplex amplicon pyrosequencing bias amplification. Appl Environ Microbiol. 2011;77(21):7846–7849. doi:10.1128/AEM.05220-11

41. Bankhead P, Loughrey MB, Fernández JA, et al. QuPath: Open source software for digital pathology image analysis. Sci Rep. 2017;7(1):1–7. doi:10.1038/s41598-017-17204-5

42. Anderson MJ. A new method for non-parametric multivariate analysis of variance. Austral Ecol. 2001;26(1):32–46. doi:10.1111/j.1442-9993.2001.01070.pp.x

