## Supplemental Table 1 for "Susceptibility to inflammatory bowel diseases promotes invasive carcinomas in a murine model of ATF6-driven colon cancer"

**Supplement Table 1: IBD cohort description**

| **Disease phenotype** | **CD (36)** | **UC (32)** |
|---|---|---|
| Gender (male/female) | (14/22) | (22/10) |
| **Disease Activity Index** | | |
| Harvey-Bradshaw Index (HBI) (0 - >16) | 9 ± 6.5 | NA |
| Crohn's Disease Endoscopic Index of Severity (CDEIS) (0 - 44) | 14.3 ± 9.2 | NA |
| Mayo score (0 - 12) | | 7.5 ± 2.3 |
| Ulcerative Colitis Endoscopic Index of Severity (UCEIS) (0 - 8) | | 4.8 ± 1.5 |
| **Smoking** | | |
| Active smoker | 8 | 1 |
| Non-smoker | 18 | 17 |
| Ex-smoker | 8 | 11 |
| NA | 2 | 3 |
| **Corticosteroids treatment** | | |
| Currently on treatment | 10 | 10 |
| No Corticosteroids pre-treatment | 4 | 3 |
| Pre-treatment with Corticosteroids | 21 | 19 |
| NA | 1 | |
| **Biological therapy administered** | | |
| 1: INFLIXIMAB | 8 | 9 |
| 2: ADALIMUMAB | 13 | 2 |
| 3: GOLIMUMAB | 0 | 9 |
| 4: USTEKINUMAB | 9 | 0 |
| 5: VEDOLIZUMAB | 5 | 9 |
| NA | 1 | 1 |
